# Single-molecule systems for detection and monitoring of plasma circulating nucleosomes and oncoproteins in Diffuse Midline Glioma

**DOI:** 10.1101/2023.11.21.568019

**Authors:** Nir Erez, Noa Furth, Vadim Fedyuk, Jack Wadden, Rayan Aittaleb, Kallen Schwark, Michael Niculcea, Madeline Miclea, Rajen Mody, Andrea Franson, Augistine Eze, Niku Nourmohammadi, Javad Nazarian, Sriram Venneti, Carl Koschmann, Efrat Shema

**Affiliations:** Department of Immunology and Regenerative Biology, Weizmann Institute of Science, Rehovot, Israel; Department of Pediatrics, University of Michigan, Ann Arbor, Michigan, USA; Center for Genetic Medicine, Children’s National Medical Center, Washington, DC, USA; Department of Pathology, University of Michigan, Ann Arbor, Michigan, USA

## Abstract

The analysis of cell-free tumor DNA (ctDNA) and proteins in the blood of cancer patients potentiates a new generation of non-invasive diagnostics and treatment monitoring approaches. However, confident detection of these tumor-originating markers is challenging, especially in the context of brain tumors, in which extremely low amounts of these analytes circulate in the patient’s plasma. Here, we applied a sensitive single-molecule technology to profile multiple histone modifications on millions of individual nucleosomes from the plasma of Diffuse Midline Glioma (DMG) patients. The system reveals epigenetic patterns that are unique to DMG, significantly differentiating this group of patients from healthy subjects or individuals diagnosed with other cancer types. We further develop a method to directly capture and quantify the tumor-originating oncoproteins, H3-K27M and mutant p53, from the plasma of children diagnosed with DMG. This single-molecule system allows for accurate molecular classification of patients, utilizing less than 1ml of liquid-biopsy material. Furthermore, we show that our simple and rapid detection strategy correlates with MRI measurements and droplet-digital PCR (ddPCR) measurements of ctDNA, highlighting the utility of this approach for non-invasive treatment monitoring of DMG patients. This work underscores the clinical potential of single-molecule-based, multi-parametric assays for DMG diagnosis and treatment monitoring.

## Introduction

Diffuse midline glioma (DMG) is a highly aggressive and fatal form of pediatric brain tumor typically arising in the midline structures of the brain, such as the thalamus, pons, cerebellum, and spinal cord ^1,2^. Diagnosis and monitoring of DMG rely heavily on Computed Tomography (CT) and Magnetic Resonance Imaging (MRI), due to the risks associated with invasive neurosurgical procedures such as routine biopsies. However, these imaging modalities are often insufficient to guide proper treatment and to monitor response. While characteristic imaging features combined with the presented symptoms allow diagnosis even without histological confirmation, recent advances in genetic technologies highlight the importance of molecular information for proper disease classification ^3,4^. Thus, despite the associated risk, biopsies are conducted on newly diagnosed cases to guide treatment decisions ^5,6^. Moreover, anatomical changes resulting from tumor progression or treatment response are hard to detect by imaging alone. MRI imaging often fails to discriminate between actual tumor progression and treatment-related pseudo-progression ^7,8^. Thus, there is a critical need to develop novel non-invasive tools for molecular-based diagnosis and monitoring of these tumors.

Detailed molecular analysis of DMG revealed a striking recurrence of lysine 27 to methionine substitution in histone H3 (H3-K27M) in more than 80% of cases. This mutation, occurring on either H3F3A or HIST1H3B/C genes, elicits widespread changes in chromatin patterns and gene expression ^9–13^. Indeed, the current classification of DMG relies on H3-K27 status, with H3-K27 altered DMG defined as a different clinical entity, showing a significant dismal prognosis compared to pediatric high-grade gliomas harboring WT H3 ^2,4^. This identification of DMG-specific hotspot driver mutations emphasizes the potential value of liquid biopsies for the diagnosis and disease monitoring of DMG.

As tumor cells die, various biomarkers, including cell-free tumor DNA (ctDNA) and proteins, are released to surrounding biofluids. Capturing and quantifying these biomarkers can provide accurate information regarding tumor burden and disease state, thus providing a minimally invasive alternative to the classical tissue biopsy ^14,15^. In the last decade, analysis of ctDNA in plasma and cerebrospinal fluid (CSF) of glioma patients has emerged as a prominent and highly useful tool for diagnostics, as well as for guiding and monitoring personalized treatment protocols ^16–18^. Importantly, liquid biopsy quantification of H3-K27M mutant alleles and additional genic biomarkers, using various strategies, can facilitate and highly improve patient surveillance and even predict treatment response, thus enhancing and simplifying glioma clinical trials ^19–23^.

While these sequencing-based approaches of ctDNA reveal rich genetic information present in CSF or plasma, they suffer from several limitations. Currently, large amounts of input material and technologically advanced instruments are required, hindering their application in large-scale studies or for screening purposes. Also, the economics of sequencing generally requires batching multiple patient samples to drive down costs, leading to long turnaround time of several weeks or even months. Furthermore, reliable analysis of ctDNA in plasma samples is still limited, requiring routine collection of CSF, a more invasive and complicated procedure ^20^. Finally, in most cases, only a single layer of information is measured (i.e. mutations analysis or DNA methylation status), limiting the output and sensitivity of these tests.

The ctDNA that circulates in the blood is packaged in the form of nucleosomes, which maintain their cell-specific histone modification patterns that are indicative of their tissue-of-origin ^24–28^. We recently established a proof-of-concept for single-molecule profiling of the Epigenetics of Plasma Isolated Nucleosomes (EPINUC) and showed its utility for detecting colorectal and pancreatic cancers ^29^. EPINUC profiles the combinatorial histone modification patterns of nucleosomes captured from plasma, in addition to providing single-molecule measurements of DNA methylation and cancer-specific protein biomarkers. This multi-modal data, obtained from less than 1ml of plasma samples taken from a cohort of ∼170 subjects, enabled highly accurate detection of cancer, even at early stages.

Here, we apply our single-molecule imaging platform to accurately detect various clinically relevant, low-abundant, biomarkers from plasma of DMG patients. Multi-modal analysis of plasma circulating nucleosomes revealed DMG-specific epigenetic patterns. We further developed a method to directly capture and quantify H3-K27M plasma circulating nucleosomes, allowing the classification of patients according to H3 status. Finally, we complement the analysis with single-molecule detection and droplet-digital PCR (ddPCR) measurements of both mutant H3 and p53 in the plasma of DMG patients during the course of chemoradiation with ONC201/206. Overall, our work establishes a novel single-molecule technology for non-invasive DMG diagnosis and patient monitoring.

## Results

### EPINUC distinguishes subjects with DMG from other cancer types and healthy individuals

We recently showed the utility of epigenetic profiling of cell-free nucleosomes (cfNuc) that circulate in the plasma to distinguish between healthy individuals and Colorectal Cancer (CRC) or Pancreatic Ductal Adenocarcinoma (PDAC) patients ^29^. To test whether this approach can provide valuable data also for brain tumors, and specifically for the aggressive DMG tumors which are difficult to access surgically due to their midline location, we applied EPINUC to plasma samples collected from DMG patients. EPINUC allows single-molecule detection of six active and repressive histone modifications on millions of individual nucleosomes directly captured from plasma (Fig. 1A and ‘Methods’). Briefly, cfNuc were tagged with a fluorophore and polyadenylated using enzymatic reactions. Labeled and tailed nucleosomes were anchored to a PEGylated surface via A-T hybridization and monitored with a panel of fluorescently tagged antibodies targeting different histone post-translational modifications (PTMs). TIRF imaging allowed for direct counting of single-nucleosomes anchored to the microscope coverslip, as well as their modification patterns. This generated a multi-parametric, quantitative, epigenetic dataset consisting of the total number of cfNuc, the percentage of nucleosomes modified with each modification, the ratio between pairs of modifications, and the percentage of nucleosomes that carry combinatorial patterns of two modifications (Fig. 1B).

**Fig. 1:**
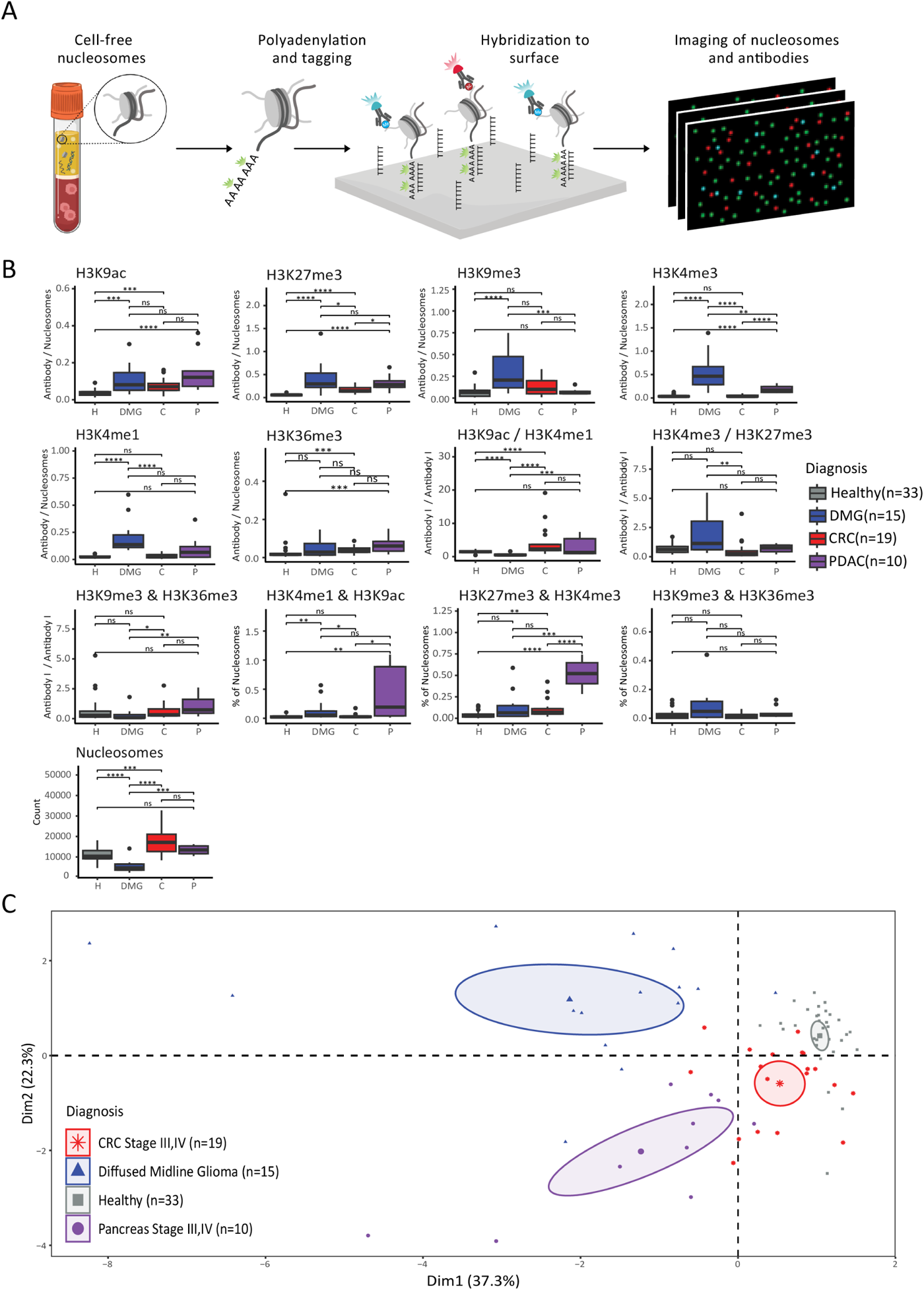
EPINUC distinguishes DMG patients from healthy individuals and subjects with colorectal or pancreatic cancer. **A.** Experimental scheme: cell-free nucleosomes (cfNuc) from plasma samples are processed in one-step reaction combining two enzymatic processes: repair of DNA ends by Klenow polymerase and addition of a poly-A tail by Terminal Transferase (TdT). The reaction contains a combination of unmodified dATPs together with fluorescently labeled dATPs (Cy3-dATP) to label nucleosomes. Next, cfNuc are anchored to PEGylated-poly-T surface via dA:dT hybridization. Anchored nucleosomes are incubated with fluorescently labeled antibodies targeting multiple histone modifications. Finally, nucleosome positions and time-lapse imaging of antibodies’ binding events are recorded by TIRF microscopy. **B.** Box plot representation of EPINUC parameters (histone PTMs levels, ratios and combinations, as indicated above the plot) measured from plasma samples collected from healthy, DMG, CRC and PDAC individuals. Box plot limits indicate 25–75% quantiles, the middle lines indicate the median, upper and lower whiskers denote the largest and smallest values, respectively, no further than 1.5× the interquartile range from the hinge and the dots represents data points outside this range. Number of samples in each group is noted in the legend. P values were calculated by Wilcoxon rank-sum exact test and adjusted by Bonferroni Correction for multiple comparisons. * P value < 0.05, ** P value < 0.01, *** P value < 0.001, ns-non significant. **C.** Principal Component Analysis (PCA) for the following EPINUC parameters: H3K4me1/Nuc, H3K9me3/Nuc, H3K9ac/Nuc, H3K27me3/Nuc, H3K9ac/H3K4me1, H3K36me3/H3K9me3, H3K27me3 & H3K4me3, and Nucleosomes count. Sample groups are color-coded as indicated in the plot; each dot represents one plasma sample. Ellipse represents 95% confidence interval for the barycenter of each group (denoted with a larger symbol).

We compared the epigenetic patterns measured in the plasma of 15 individuals diagnosed with DMG, to those previously measured in late-stage CRC patients (stages III, IV), PDAC patients as well as healthy subjects ^29^. This analysis revealed multiple epigenetic parameters that differed significantly between DMG samples and healthy controls, stressing the power of EPINUC in detecting tumor-associated epigenetic alterations (Fig. 1B). For example, plasma of DMG patients showed increased levels of H3K9ac-, H3K4me1-, H3K9me3-, and H3K4me3-modified nucleosomes, while presenting lower H3K9ac to H3K4me1 ratio, compared to healthy individuals. The higher number of nucleosomes marked by H3K9ac and H3K9me3s is in agreement with previous studies showing upregulation of these marks in DMG tumors ^13,30,31^. MLL1, the enzymatic complex that deposits H3K4me3, was also reported to be deregulated in H3-K27M-mutant DMGs, and indeed H3K4me3 levels were higher in these patients compared to both healthy individuals and subjects with CRC/PDAC ^13,30^. Yet, while these data largely agree with previous reports indicating global elevation of these marks in pediatric high-grade gliomas, it is important to note that alterations in the epigenetic patterns of circulating nucleosomes do not necessarily originate from the contribution of tumor cells, and may also be attributed to changes in the tumor’s microenvironment and the immune system ^24^.

In contrast to other cancers, the plasma of DMG patients did not show elevated number of cfNucleosomes, highlighting the sensitivity of our assay in detecting epigenetic changes even for brain tumors that are likely not shedding a large number of nucleosomes to the blood. Surprisingly, we also observed an increase in H3K27me3-modified nucleosomes in DMG patients, despite the loss of this modification in DMG tumors ^11,32,33^. As noted above, this increase may stem from the systemic response to the tumor, or a response to different anti-cancer treatments (such as ONC201^34^) that these patients received. Importantly, H3K27me3 levels were also elevated in plasma taken from CRC and PDAC patients, suggesting a pan-cancer phenomenon that should be further explored in additional systems. Overall, these data point to the advantages of EPINUC in integrating data from multiple high-resolution epigenetic measurements that may originate from the tumor cells or the tumor’s microenvironment.

Next, we performed principal-component analysis (PCA) to visualize the distribution of DMG samples across the epigenetic parameters that showed most variability between all groups. The PCA showed a distinct spatial separation between all groups, with DMG samples forming a separate cluster, in agreement with the unique epigenetic patterns found in DMG plasma samples (Fig. 1C). This analysis highlights EPINUC’s ability to differentiate between multiple types of cancers solely based on measurements of a plethora of epigenetic parameters present in plasma.

### Enrichment and detection of plasma circulating H3-K27M nucleosomes by single-molecule imaging

Determining H3 status is key for proper diagnosis and classification of pediatric gliomas, which currently require staining or sequence analysis of tumor biopsy ^4^. While our EPINUC platform could clearly distinguish DMG patients from subjects with CRC, PDAC or healthy controls, we did not detect a clear separation between H3-K27M-mutant and WT DMG samples (Fig. S1). This result agrees with the notion that most epigenetic alterations in the plasma of cancer patients represent the distinct tissue-of-origin contribution (for example, colon for CRC patients), rather than cancer-specific alterations ^35–37^ .

To tackle the challenge of specific detection of the H3-K27M mutation in DMG patients, we aimed to develop a simple method to directly probe these mutant nucleosomes in plasma, thus greatly improving the diagnostic power of our liquid biopsy tests. We reasoned that while H3-K27M nucleosomes, originating from tumor cells, may exist in plasma, they are likely present in very low levels, making single-molecule imaging ideal for their measurement. Thus, we developed a novel strategy for on-surface enrichment of plasma circulating H3-K27M nucleosomes. We modified surface chemistry to contain PEG–streptavidin coating, which allows anchoring of biotin-conjugated antibodies specific for the H3-K27M-mutant histone ^30^ (see ‘Methods’). These antibodies can then specifically capture fluorescently tagged H3-K27M-mutant nucleosomes from plasma, allowing imaging by TIRF microscopy. Importantly, WT nucleosomes will not bind tightly to the K27M-specific antibody, thus generating a surface enriched with H3-K27M-mutant nucleosomes (Fig. 2A).

**Fig. 2:**
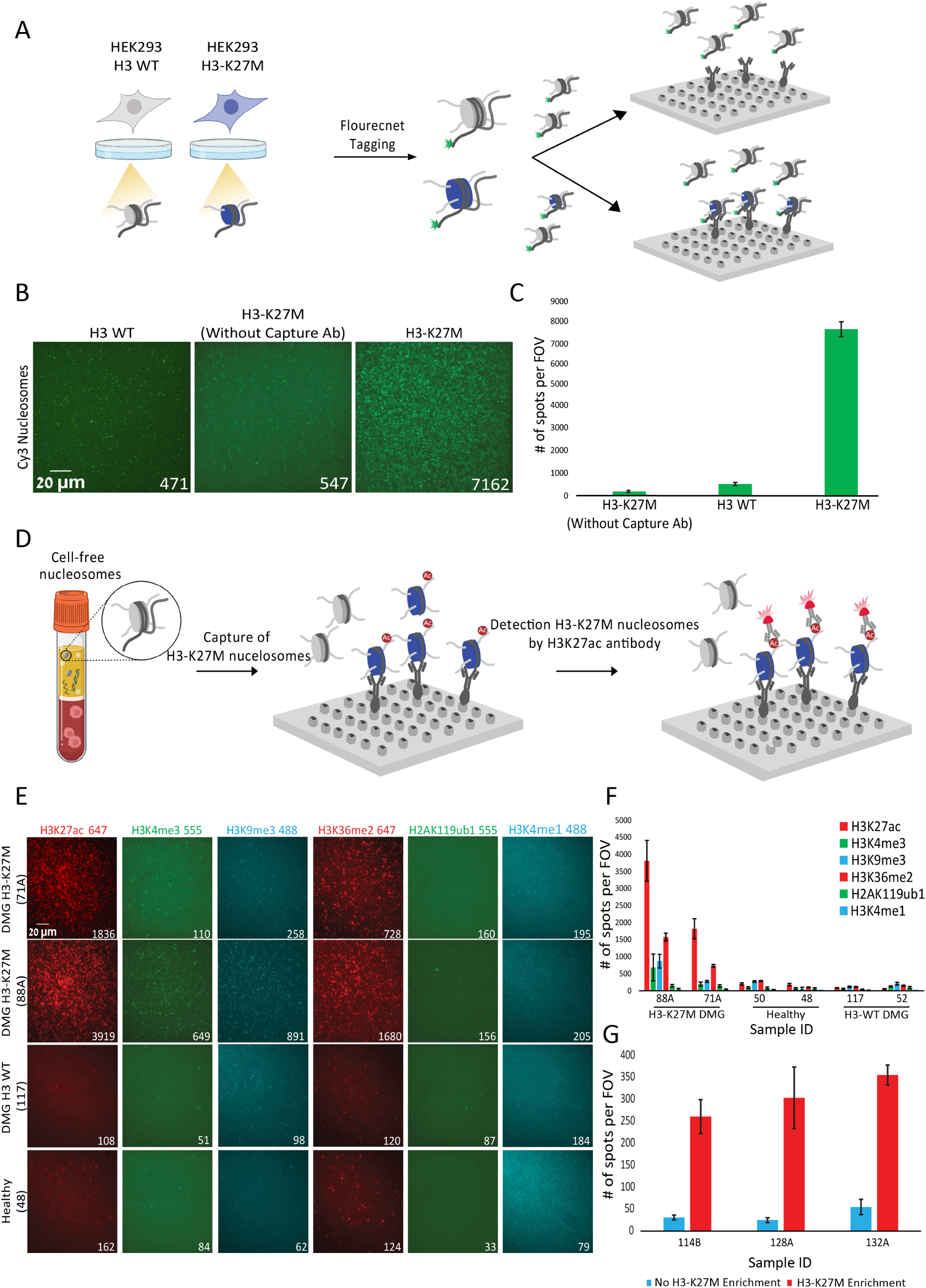
Single-molecule approach for enrichment of plasma circulating H3-K27M-mutant Nucleosomes. **A.** Experimental scheme: nucleosomes are extracted from HEK293 cells ectopically expressing either WT or mutant H3 and the DNA is fluorescently labeled with Cy3 by Klenow polymerase and Cy3-dATP. Next, labeled H3-K27M nucleosomes are captured on PEG–streptavidin surface covered with biotinylated antibodies targeting the H3-K27M mutant histone (α-H3-K27M) and imaged by TIRF imaging. Nucleosomes containing WT H3 do not bind the microscope coverslip and are washed prior to imaging. **B**. Representative TIRF images of Cy3 signal, corresponding to surface bound nucleosomes, extracted from HEK293 cells expressing WT or mutant histone. Nucleosomes extracted from cells expressing H3-K27M were loaded also on a surface absent of α-H3-K27M antibodies (middle panel), to demonstrate antibody-dependent capture. Numbers within images represent counted spots; each spot correspond to a single surface bound nucleosome. Nucleosomes extracted from cells expressing mutant H3 show higher binding to surface coated with α-H3-K27M antibodies. **C**. Quantification of the experiment described in B. Data is presented as the mean +/-s.d. of 50 fields of view (FOVs) per sample. **D**. Experimental scheme: cfNuc from plasma are immobilized on a PEG–streptavidin covered with biotinylated α-H3-K27M antibody. Captured H3-K27M nucleosomes are detected by incubation with fluorescently-labeled antibody followed by TIRF imaging. **E**. Representative TIRF images of different antibodies targeting the indicated histone PTMs, while incubated on surfaces enriched for plasma circulating H3-K27M nucleosomes (as described in D). Antibodies were tested on plasma samples from healthy individuals, as well as DMG patients carrying WT H3 or H3-K27M. Numbers within images represent counted spots; each spot correspond to a single surface bound nucleosome. H3K27ac and H3K36me2 antibodies are suitable for detection of surface bound H3-K27M nucleosomes. **F-G**. Quantification of the single-molecule signal obtained for H3K27Ac antibody when incubated with and without H3-K27M enriched cfNuc from different plasma samples. Data is presented as the mean +/-s.d. of 50 FOVs per sample.

To test this approach, we established HEK293 cells ectopically expressing the mutant H3-K27M-histone or WT H3 as control ^30^. Nucleosomes from these cells were extracted and labeled with Cy3 (green), then added to the flowcell containing a surface coated with α-H3-K27M antibody as described above, followed by imaging (Fig. 2B-C). Indeed, we observed a significant increase (∼10-fold) in signal for the sample expressing H3-K27M, compared to nucleosomes from cells expressing WT H3. This validated the preferential binding and enrichment of mutant nucleosomes on the surface. To further validate that this enrichment is dependent on the α-H3-K27M antibody, we incubated nucleosomes extracted from H3-K27M mutant cells on a PEG-streptavidin surface without α-H3-K27M antibody. As expected, this resulted in background level signal, comparable to the one obtained for nucleosomes extracted from control cells (WT H3). This data confirmed our ability to selectively bind and enrich H3-K27M-mutant nucleosomes on the microscope surface.

With the above-described enrichment methodology in hand, we set to probe H3-K27M-mutant nucleosomes directly from plasma. In comparison to cell extracts, plasma samples are expected to contain very low levels of mutant nucleosomes, and we reasoned that direct labeling of total cfNuc may not be efficient enough to capture such rare events. In addition, we aimed to minimize sample processing, to avoid potential loss of nucleosomes and ensure maximal detection rate. Thus, instead of enzymatically labeling nucleosomes in plasma, we explored an alternative strategy, using fluorescently-labeled antibodies to visualize the H3-K27M-mutant nucleosomes that are captured on the surface (Fig. 2D). Antibodies targeting the unmodified core histones (i.e., antibodies for total H3/H4/H2A/H2B) show very low signals in our single-molecule system; this is presumably due to the high evolutionary conservation of histones, low accessibility for antibody binding at the globular domain, or potential binding-interference by PTMs when the antibody is targeted to the histone tail. Thus, we screened six antibodies targeting different histone PTMs for their ability to detect surface-bound plasma circulating H3-K27M nucleosomes (Fig. 2E). The antibodies targeting H3K27ac and H3K36me2 modifications reliably detected surface-bound mutant nucleosomes, showing high difference between H3-K27M DMG samples compared to samples taken from H3 WT DMG patients and healthy individuals (Fig. 2E-F and S2A). Indeed, both modifications have been reported to be enriched on H3-K27M nucleosomes, supporting their use for detection of surface-bound H3-K27M nucleosomes ^30,38,39^. H3K27ac signal showed a broader dynamic range across the samples tested, rendering it a better candidate for our assay. Of note, H3K27ac signal on H3-K27M enriched surfaces was much higher compared to H3K27ac global levels in each sample, as quantified using the standard EPINUC protocol (Fig. 2G and S2B). This further highlights the importance of the “on-surface” enrichment step for the detection of low abundant mutant nucleosomes.

### Single-molecule detection of H3-K27M nucleosomes differentiates between H3-K27M and H3 WT DMG patients

To validate our H3-K27M-H3K27ac single-molecule detection system, we applied it to a cohort comprising of 21 plasma samples from healthy adults, 9 samples from healthy children, 46 samples from H3-K27M DMG patients, and 10 samples from H3 WT DMG patients (Table S3). The DMG cohort consists of samples from individuals across treatment stages. Notably, single-molecule counting of H3K27ac, corresponding to H3-K27M surface-bound nucleosomes, was significantly higher in H3-K27M DMG samples (Fig. 3A-B). All other tested groups (DMG patients carrying WT H3, healthy adults and healthy children) showed similar low signal for H3-K27M-H3K27ac, due to the absence of H3-K27M mutation in their genome (thus, mutant nucleosomes are not captured on surface). Of note, a small number of individuals diagnosed with H3-K27M DMG showed low signals, potentially reflecting smaller tumors or tumor remission due to successful response to treatment at the time of sample collection.

**Fig. 3:**
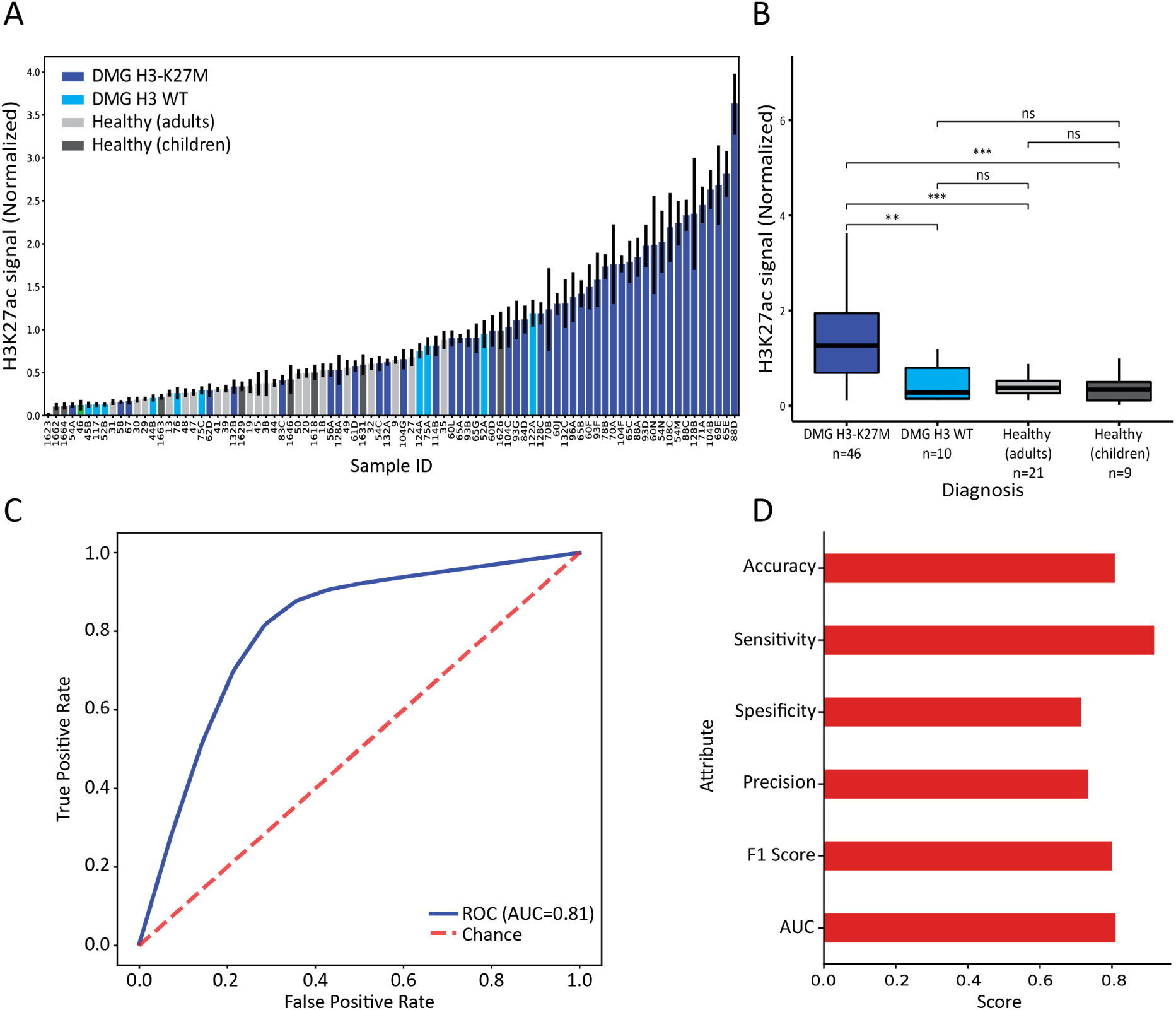
Single-molecule detection of mutant nucleosomes in plasma differentiates H3-K27M from H3 WT DMG patients. **A**. Plasma samples of DMG patients carrying WT H3 or H3-K27M-mutant (light blue and dark blue, respectively) and plasma of healthy adults or children (light grey and dark grey, respectively), were analyzed as described in Fig. 2D. Mutant nucleosomes captured on surface were detected by H3K27ac antibody. Each bar represents a subject and data is presented as the mean of the normalized H3K27ac counts +/-s.d. of 50 FOVs per sample. **B**. Box plot representation of the data in A. n denotes the number of samples in each group. Box plot limits indicate 25–75% quantiles, the middle lines indicate the median and the upper and lower whiskers denote the largest and smallest values, respectively, no further than 1.5× the interquartile range from the hinge. P values were calculated by Wilcoxon rank-sum exact test and adjusted by Bonferroni Correction for multiple comparisons. ** P value < 0.01, *** P value < 0.001, ns-non significant. **C**. ROC curve discriminates between plasma samples containing WT H3 nucleosomes (n=40) and H3-K27M-mutant nucleosomes (n=46) using a Naïve Bayes model. The area under the curve (AUC) for H3-K27M-H3K27ac normalized counts (Blue line), while the red dashed diagonal line indicates expected curve for random classification. **D.** H3-K27M-H3K27ac classifier performance; each bar represent the mean value of repeated (n=10,000) fourfold cross-validation across all samples.

The high H3-K27M-H3K27ac signal detected in H3-K27M DMG samples prompted us to apply machine learning algorithms in order to generate a classifier based on our single-molecule data (Fig. 3C-D and Fig. S3). The best predictive model displayed high diagnostic potential by generating a 0.81 area under the curve (AUC; 95% confidence interval (CI) of 67.2-92.9) and sensitivity of 90% (95% CI of 71.2–100) at 71% specificity (95% CI of 51.7–92.8) and 73% precision (95% CI of 59.3–88.9). While these results need to be extended to larger cohorts, they clearly highlight the potential applicability of this methodology for diagnosis and classification of DMG, in a non-invasive and cost-effective manner. As plasma sample does not undergo any processing, the test is highly simple and rapid: ∼60 minutes to obtain a robust detection.

### Detection of plasma circulating mutant p53 in DMG patients

Our ability to detect non-secreted, low-abundant, tumor-specific proteins encouraged us to explore whether our capture-detection approach can be applied to additional oncoproteins originating from glioma cells. TP53 gene, encoding the well-studied tumor-suppressor protein p53, is frequently mutated in DMG ^12^. We recently demonstrated the use of multiplexed mutant p53 detection method in plasma of CRC patients ^29^. To this end, we coated PEG-streptavidin surfaces with biotin conjugated α-p53 antibodies to allow capturing of plasma circulating p53. We then applied concurrent detection with two discrete fluorescently labeled antibodies; one antibody targeting both the wild-type and mutant p53 (’Total p53’, labeled with red) and a second antibody favoring the mutant p53 protein (’Mutant p53’, labeled with green, Fig. 4A-B).

**Fig. 4:**
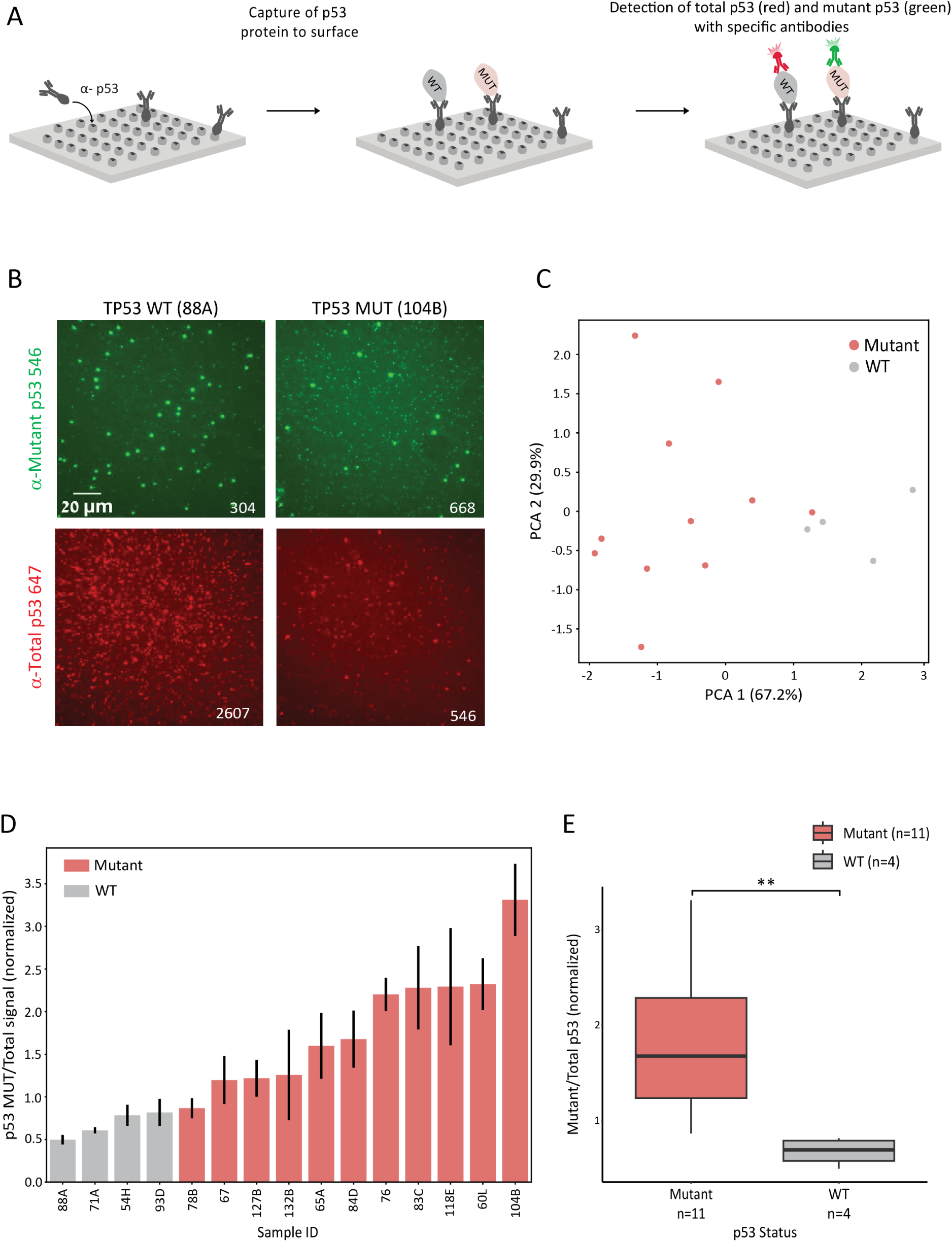
Detection of plasma circulating mutant p53 in DMG patients. **A**. Experimental scheme: biotinylated capture antibodies targeting total p53 protein (the WT and mutant form) are anchored to a PEG-streptavidin surface (α-p53). Plasma circulating p53 protein molecules are captured on surface, followed by incubation with two distinct fluorescently labeled p53 antibodies: an antibody targeting all forms p53 (’total p53’, red) and an antibody specific to the mutant conformation of p53 (’mutant p53’, green). **B.** Representative TIRF images of the indicated p53 antibodies incubated on surfaces enriched for plasma circulating p53 proteins. **C**. A cohort of 16 plasma samples of H3-K27M DMG patients was analyzed as described in A (12 samples harbor TP53 mutations and 4 harbor WT TP53). Principal Component Analysis (PCA) with the following input parameters: normalized counts of total p53 and mutant p53, and the ratio between mutant and total p53. Sample groups are color-coded according to known TP53 status; each dot represents one plasma sample. **D**. The ratio between mutant and total p53 signal for each sample is shown. Each bar represents a sample, color-coded according to known TP53 status. Data is presented as the mean +/-s.d. of 50 FOVs per sample. **E.** Box plot representation of the data in D. n denotes the number of samples in each group. Box plot limits indicate 25–75% quantiles, the middle lines indicate the median and the upper and lower whiskers denote the largest and smallest values, respectively, no further than 1.5× the interquartile range from the hinge. P values were calculated by Wilcoxon rank-sum exact test. ** P value < 0.01.

We applied this approach to 15 H3-K27M mutant DMG plasma samples (Table S3); 11 samples from patients harboring a mutation in the TP53 gene and 4 samples from patients harboring WT p53 (based on sequencing of tumor biopsies). For each sample we quantified several parameters: the signal obtained from the total-p53 antibody, the signal obtained from the mutant-specific antibody and the ratio between the two antibodies, reflecting the fraction of the mutant out of the total. Strikingly, PCA of these parameters revealed separation between the two sample groups (WT vs. MUT p53, Fig. 4C). While samples from DMG patients carrying WT p53 clustered relatively tightly together, samples from patients harboring TP53 mutation showed greater variability, possibly due to variation in the type of p53 mutation between tumors, causing slight changes in the performance of the mutant-specific antibody. Of note, the ratio between signals of the two antibodies (mutant-specific vs. total) is the most significant parameter contributing for the separation between the groups (Fig. 4D-E). These results suggest specific and sensitive detection of mutant p53 in the plasma of DMG patients that carry this mutation.

### Multiplexed and multimodal analysis of serial plasma samples may inform DMG treatment response in patients treated with ONC201/206

Prior work has shown that fluctuations in ctDNA present in patients’ plasma and CSF predict tumor progression, and could potentially be used to augment magnetic resonance imaging (MRI) for treatment response monitoring ^19^. Specifically, decrease in ctDNA levels, resulting from reduction in plasma circulating tumor cell debris, was shown to correlate with clinical improvement. We hypothesized that mutant protein fluctuations, which can be measured by our single-molecule systems in a rapid, simple, and cost-effective manner, might have similar utility in treatment response monitoring, and provide additional important information to help clinicians best monitor and manage the disease. To test this hypothesis, we retrospectively performed multimodal (protein and ctDNA) and multiplexed (H3.3-K27M and TP53) analysis of serial samples from three H3.3-K27M and TP53 mutant DMG patients: UMPED128, and UMPED132, and UMPED65, undergoing chemoradiation treatment with ONC201 or ONC206. ONC201/206 are two promising small-molecule imipridones, currently in clinical trials for DMG patients ^40^. In addition to single-molecule measurements of tumor oncoproteins, we conducted ddPCR analysis of ctDNA, quantifying H3.3K27M and the patient specific TP53 mutation, over different timepoints during treatment.

UMPED128 (Fig. 5A-B) is a 9-year-old male diagnosed in February 2022 with DMG (H3.3-K27M and TP53 R273C), showing multifocal involvement. He was given radiation and temozolomide before starting ONC206 at 4 months after diagnosis, progressing 6 months later. Both H3.3-K27M and TP53R273C ctDNA levels initially decreased, matching radiographic response before trending slightly upwards at early clinical progression (Fig. 5B). Strikingly, the single-molecule H3-K27M-H3K27ac measurements matched tumor area, showing an initial decrease due to treatment, followed by a robust increase upon disease progression. This robust increase in H3-K27M-H3K27ac nucleosomes matched the large increase in tumor size and vascularized necrosis notable in the final MRI taken at progression. Notably, the fraction of mutant p53 protein (relative to the WT-p53) trended upwards immediately after treatment and continued an upward trend through treatment and progression, perhaps indicating the lack of effective response to treatment.

**Fig. 5:**
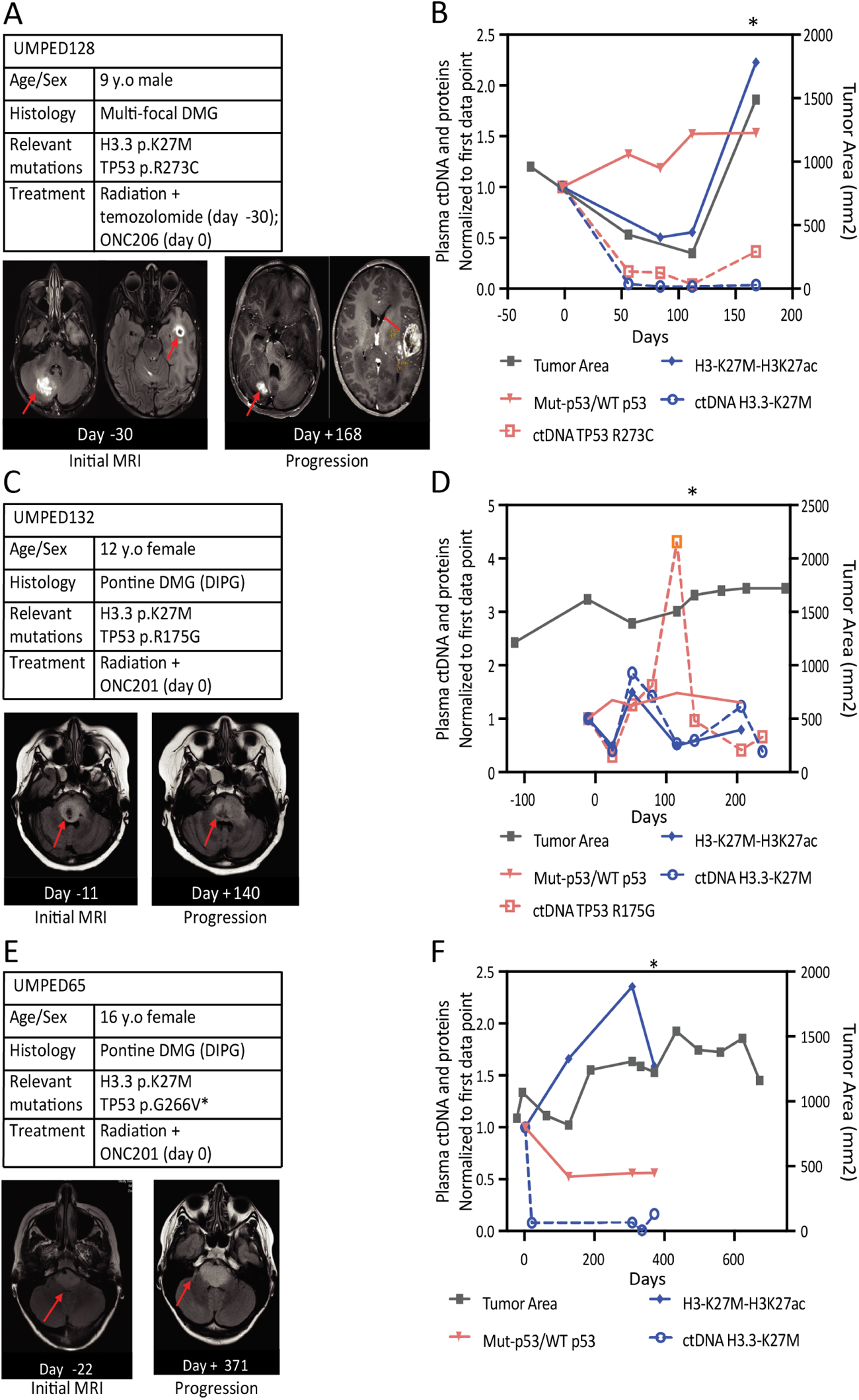
Multi-modal and multiplexed analysis of serial plasma samples from patients undergoing ONC201 treatment. Clinical features, initial MRI, MRI of disease at clinical progression (A, C, E), and results from multimodal and multiplex H3/TP53 assays performed on serial timepoint plasma samples plotted along with tumor cross sectional area (B, D, F) for three patients: **A-B.** UMPED128, **B-C.** UMPED132 and **C-D.** UMPED65. For UMPED65, TP53 p.G266V ddPCR assay was not performed due to assay malfunction. Day-0 corresponds to initiation of treatment with ONC201/206. An asterisk indicates diagnosis with progressive disease.

UMPED132 (Fig. 5C-D) is a 12-year-old female diagnosed in March 2022 with DMG (H3.3-K27M and TP53R175G). She received radiation and started ONC201 at 4 months, progressing 6 months later. Similar to UMPED128, ctDNA signals of both H3.3-K27M and TP53 initially decreased after treatment with ONC201 along with a noticeable decrease in tumor area (Fig. 5D). Importantly, also in this case the levels of H3K27M-H3K27ac nucleosomes measured in plasma matched the trends of ctDNA signals, showing first a decrease in response to treatment, followed by an increase upon disease progression. As in patient UMPED128, mutant p53 protein ratio immediately trended upwards after treatment with ONC201 and before radiographic progression. Of note, just prior to clinical progression, TP53R175G ctDNA signal increased dramatically along with a slight increase in mutant p53 ratio, while H3.3-K27M ctDNA and nucleosomes actually decreased. This divergence may stem from an increase in tumor heterogeneity or overall complexity under treatment, and highlights the importance of multimodal and multiplex monitoring.

UMPED65 (Fig. 5E-F) is a 16-year-old female diagnosed in December 2018 with DMG (H3.3-K27M and TP53R266V). Two months after diagnosis, she began taking ONC201 with concurrent radiation, progressing 12 months later. At this point, she remained on ONC201 and was started concurrently on the HDAC inhibitor panobinostat and later on the PI3K inhibitor paxalisib. After initial treatment, a significant drop in H3.3-K27M ctDNA signal was noted, correlating with clinical improvement. Unfortunately, we did not have plasma sample for single-molecule measurements of the mutant nucleosomes at this early time point after treatment. Interestingly, despite initial reduction in tumor area following treatment, between days 124 and 306 there is a notable increase in tumor size, which matches a continuous increase in the levels of mutant H3-K27M-H3K27ac detected in the plasma. Of note, ctDNA levels throughout this time window showed only a very mild increase. Strikingly, at time point 306 days there is a mild but consistent reduction in tumor size, which matches a reduction in ctDNA levels as well as H3-K27M-H3K27ac mutant nucleosomes. Unfortunately, following this time point the disease of this patient progressed, and we could not obtain additional plasma samples. An initial indication for disease progression may be seen in the increase in ctDNA between the last two measurements. Overall, out of all measurements (ctDNA, p53 ratio and mutant nucleosomes) in the three patients across the treatment, the trends observed for the single-molecule measurements of H3-K27M-K27ac showed the best match to tumor area (MRI). Although limited to a small number of samples, these results serve as a proof-of-concept for the clinical potential of our single-molecule imaging platform to non-invasively detect and monitor the levels of mutant p53 and H3-K27M mutant nucleosomes in plasma of DMG patients. Larger studies will be required to further improve and verify these early observations.

## Discussion

Developing simple assays which can reliably detect and quantify multiple analytes in the blood is crucial for clinical implementation of liquid biopsy approaches. This task is highly challenging, as most clinically-relevant analytes are present at very low concentrations in plasma, and input sample is limited. Recent works established the use of a single epigenetic modification; DNA methylation, as a promising direction for blood-based multi-cancer detection ^41,42^. Additional epigenetic features, such as the fragmentation patterns of cfDNA, may also be highly relevant for liquid-biopsy analysis of brain tumors, and specifically in pediatric cases ^43,44^. Yet, despite the clear advantage of liquid-biopsies over tissue sampling, detection of tumor-derived cfDNA in gliomas has been difficult ^43^. We show that multi-parametric analysis of plasma circulating nucleosomes allows to distinguish between healthy individuals and DMG patients. Moreover, the multiple epigenetic parameters captured with our method differed between DMG patients and other cancer types such as CRC or PDAC, underscoring the ability of these features to reveal the tumor type. Of note, assaying multiple parameters is likely to reduce confounding factors such as age, treatment, etc. Yet, some differences between the groups may contribute to the observed epigenetic segregation. Importantly, when assaying specifically for H3 status, by enriching for H3-K27M nucleosomes, we also controlled for age differences by analyzing plasma collected from children not-diagnosed with cancer.

While the basic EPINUC features were enough to differentiate plasma samples from DMG patients, specific and accurate classification of the H3 status of these tumors required further, case-specific tailoring of the method. Taking advantage of the high sensitivity and quantitative nature of the single-molecule imaging system, we established a novel approach for specific detection of plasma circulating H3-K27M mutant nucleosomes. We apply “on-surface” enrichment of H3-K27M nucleosomes, allowing for highly sensitive detection, without the need of any prior sample processing. This enabled accurate classification of DMG patients according to H3 mutation status. While the classical enzyme-linked immunosorbent assay (ELISA) is frequently used for detection of protein biomarkers, it has low dynamic range and is highly limited to proteins present at high concentration in plasma ^45,46^. Mass spectrometry-based methods for analysis of proteins in plasma are also biased towards more abundant proteins ^47^. We provide the first demonstration of protein-based detection of circulating H3-K27M nucleosomes in plasma, which are derived directly from brain tumors, and to the best of our knowledge cannot be detected with other methods.

The very low amount of sample needed for this enrichment analysis (∼100ul of plasma) allows multiplexing with additional tests to provide a comprehensive dataset for each patient. For example, here, in addition to H3-K27M detection, we also conducted histone modification profiling (EPINUC), as well as mutant p53 detection; all these assays combined required less than 1ml of plasma. Thus, this type of analysis can be routinely combined with additional, multimodal assays that require larger plasma volumes, such as ctDNA analysis, where >1ml of plasma is generally required to recover enough ctDNA to precisely resolve small variant allele fractions (i.e. <0.1%). H3-K27M nucleosomes and the mutant form of p53 are both non-secreted proteins that originate directly from the glioma cells. Detection of these oncoproteins in the plasma confirms the penetrance of these tumor-derived proteins across the blood-brain barrier and highlights the sensitivity of our method ^48^. Importantly, this approach can be easily adapted for detection of additional proteins, which may further improve the accuracy of DMG classification, overcoming inter-patient heterogeneity. Furthermore, our study lays the framework for detecting a variety of driving oncoproteins, such as EZHIP, KRAS, and BRAF, in the context of pediatric and adult cancers.

As most DMG patients enroll in clinical trials and are monitored routinely, it is of high importance to characterize and establish reliable biomarkers of response. Currently, CSF is understood to be a superior bio-fluid for disease monitoring ^19,23^ because it contains much higher concentrations of tumor-derived molecules resulting in higher-sensitivity of detection and higher-precision measurements ^16^. While lumbar punctures are minimally invasive, it is still a relatively major procedure requiring general anesthesia in the pediatric use-case. Thus, detection of tumor-originating molecules in plasma of glioma patients will highly simplify patient monitoring and reduce patient and family burden. We show, on a limited number of samples, the potential use of serial measurements of mutant p53 and H3-K27M nucleosomes in plasma. Importantly, our single-molecule measurements of H3-K27M-H3K27ac nucleosomes largely agree with tumor area and ctDNA analysis, sometimes even showing clearer and more robust trends. Interestingly, mutant p53 measurements diverged from H3-K27M-H3K27ac nucleosomes and ctDNA; regardless of ctDNA increase or decrease upon initiation of treatment, immediate increases in mutant p53 signal, and decreases in H3-K27M nucleosomes after treatment correlated with decrease in progression free survival (UMPED128 and UMPED132) while opposite trends correlated with prolonged progression free survival (UMPED65). This divergence in patterns between mutant nucleosomes and mutant p53 may stem from differences in stability or secretion mode. To the best of our knowledge, this work is the first to monitor both mutant protein biomarkers and mutant ctDNA, in parallel to MRI measurements of tumor area, in serial patient samples. Future larger prospective studies will be required to validate these early findings.

Overall, we provide a proof-of-principle for the use of single-molecule based assays for diagnosis and monitoring of DMG patients. The unique and varied kinetics of each biomarker across this small cohort motivates monitoring of as many tumor-derived protein signals and ctDNA targets as possible to increase the utility of liquid biopsies for treatment response monitoring.

## Acknowledgments

E.S. is an incumbent of the Lisa and Jeffrey Aronin Family Career Development chair. This research was supported by grants from the European Research Council (ERC801655, ERC_PoC_963863), Emerson Collective, The Israeli Science Foundation (1881/19), the Israel Cancer Research Fund – Research Career Development Award and the Weizmann – Swiss Society Institute for Cancer Prevention Research.

C.K. is supported by NIH/NINDS Grant R01-NS124607 and R01-NS119231 and Department of Defense Grant CA201129P1, the University of Michigan Chad Carr Pediatric Brain Tumor Center, the ChadTough Defeat DIPG Foundation, the DIPG Collaborative, Catching Up With Jack, The Pediatric Brain Tumor Foundation, The Yuvaan Tiwari Memorial Foundation, The Morgan Behen Golf Classic, and the Michael Miller Memorial Foundation. J.W. is supported by the National Institutes of Health under award number T32HL749, the University of Michigan Chad Carr Pediatric Brain Tumor Center, Catching Up With Jack, and the Pediatric Brain Tumor Foundation.

## Author contributions

N.E., N.F. and E.S. designed the study and wrote the manuscript. C.K. and S.V. assisted with designing experiments and writing the manuscript. N.E., N.F. and V.F. performed the experiments and analyzed the data. J.W. and R.A. performed DNA extractions and designed, performed, and analyzed all ddPCR-based experiments. K.S., M.N., M.M., R.M., A.F., A.E., J.N. and C.K. assisted with the collection of plasma samples from healthy children and DMG patients.

## Data and materials availability

All data are available in the main text or the supplementary materials.

## Materials and Methods

### Participants

All clinical studies were approved by the Israeli Ministry of Health Ethics Committees (Helsinki applications 091-2020 and 0198-14-HMO). Informed consent was obtained from all individuals before blood sampling.^37^

For DMG plasma samples, informed consent/assent was obtained on the IRB-Approved protocol Sample Repository for Pediatric Hematology/Oncology (HUM00123426).

For the healthy patient samples, specimens were obtained on the Children’s Nationla Hospital IRB-approved protocol Biomarker Identification In Pediatric Brain Tumors (Pro00000747).

### Plasma collection

Blood samples were collected in Vacuette K3 EDTA tubes and transferred immediately to ice. Next, blood was centrifuged (10 min at 1,500g and 4 °C), and the supernatant was transferred to a fresh 50-ml tube and centrifuged again (10 min at 3,000g and 4 °C). The supernatant was collected (while avoiding the buffy coat layer) and used as plasma for all experiments. Plasma was analyzed fresh or flash-frozen and stored at −80 °C for future analysis.

UM-derived samples: blood samples were collected in BD Vacutainer® K2 EDTA tubes and transported at room temperature back to the lab. Next, blood was centrifuged (15 min at 2000 g and 4 °C). Then, the supernatant was collected (while avoiding the buffy coat layer) and stored at −80 °C for future analysis.

### DNA extraction and pre-amplification

ddPCR primers, probes, and protocols were designed in house or via Biorad or Integrated DNA Technologies design services. Assays were validated in singleplex and adjusted to improve separation of wildtype and mutant droplet populations. Patient cell-free DNA was extracted from 1ml-1.2ml of patient plasma using the QIAamp Circulating Nucleic Acid Kit (QIAGEN # 55114) according to the manufacturer’s instructions. In all cases, multiple elutions were performed to extract as much DNA as possible from each sample. Eluted DNA was then vacuum concentrated at 45 °C (Thermo Scientific DNA130) and used in full as input for pre-amplification. To enable multiplex ddPCR analysis from limited patient plasma, extracted DNA was pre-amplified in duplex using a combination of H3.3 and TP53 ddPCR primer sets according to known patient mutations. Duplex PCR was performed using Q5 Hotstart 2x Master Mix (NEB) with final primer concentrations of 0.5uM. Duplex product was then quantified using Qubit 3.0 Fluorometer (Invitrogen; dsDNA HS assay) and diluted to approximately 100,000 copies/ul before being used as input to singleplex H3.3 K27M or TP53 mutant ddPCR. A detailed description of all primers, and protocols is described in the Supplementary Material.

### Droplet digital PCR (ddPCR) assay and analysis

A detailed description of all primers, probes, and full protocols is provided in the Supplementary Material. In brief, ddPCR reaction mixtures were prepared using ddPCR Supermix for Probes (no dUTP) (BioRad). ddPCR was performed using a BioRad QX200 droplet digital PCR system. All samples were run multiple times to adjust input copies/ul to improve assay sensitivity. Final results were gathered using 3-6 replicates. Variant allele fractions for each sample were calculated by isolating distinct populations of droplets using QuantaSoft Analysis Pro (v1.0596). Allele fractions from each replicate were averaged and normalized relative to the first data point. Primer, probe, pre-amplification, and ddPCR analysis details are provided in the Supplementary material.

### Circulating free nucleosome (cfNuc) preparation for single-molecule imaging

Plasma nucleosomes were processed as described in Fedyuk et al ^29^. Briefly, 20 µl of plasma were incubated at 37 °C for 1 h with the following reaction mixture: 10 µl of 10× green buffer (Enzymatics, B0120), 416 µM CoCl2 (Enzymatics, B0220), 1:60 protease inhibitor cocktail (PI; Sigma, P8340), 83.3 nM fluorescently labeled dATP (Jena Bioscience, NU-1611-Cy3), 83.3 µM dATP (Thermo Fisher Scientific, R0181), 5 µl of Klenow Fragment (3′→5′ exo-, NEB, M0212S), 3 µl of T4 polynucleotide kinase (NEB, M0201L) and 4 µl of terminal deoxynucleotidyl transferase (Enzymatics, P7070L). Following incubation, samples were inactivated by immediate transfer to ice.

### Cellular nucleosome preparation for single-molecule imaging

Extraction and labeling of cellular nucleosomes were performed as describes in Furth et al ^30^. Briefly, cells were collected and washed with PBS supplemented with protease inhibitors cocktail, 1:100, Sigma P8340), and HDAC inhibitors (20mM Sodium butyrate, Sigma 303410 and 10uM Vorinostat, LC Laboratories V-8477), followed by centrifugation at 3000 rpm for 3 min. Cell pellet was resuspended in 1mL of 0.05% IGEPAL (Sigma I8896) diluted in PBS (supplemented with inhibitors), centrifuged once again at 3000 rpm for 3 min. Pellet was then resuspended in Lysis buffer (100mM Tris-HCl pH 7.5, 300mM NaCl, 2% Triton® X-100, 0.2% sodium deoxycholate, 10mM CaCl2) supplemented with inhibitors and Micrococcal Nuclease (MNase, ThermoFisher Scientific, 88216). The suspension was incubated at 37°C for a time period appropriate for mono-nucleosome digestion (determined for each MNase batch), and then inactivated by addition of EGTA at a final concentration of 20mM. Next, the lysate was centrifuged for 10 min at max speed and supernatant was collected to a new tube. 10% of the lysate was used to extract DNA by AMPure SPRI beads (Beckman Coulter, A63881) and chromatin digestion was verified by resolving on 2% Agarose gel. Nucleosomes were then concentrated on an Amicon ultra-4 (Millipore, UFC810024) for 20 min followed by 1 min centrifugation at 16,000 g. Inhibitors were supplemented following concentration. For nucleosomes labeling reaction mix contained: NEBuffer™ 2 (NEB B7202), protease and HDAC inhibitors (as detailed above), 0.25 mM MnCl2, 33uM fluorescently labeled dATP (Jena Bioscience, NU-1611-Cy3),1.5ul of Klenow Fragment (3’→5′ exo-, NEB, M0212S) and 1.5ul of T4 Polynucleotide Kinase (NEB, M0201L). Sample was incubated at 37°C for 1 h and then inactivated by addition of EDTA at a final concentration of 20mM. Nucleosomes were then purified on Performa Spin Columns (EdgeBio, 13266) followed by the addition of inhibitors.

### Surface preparation for single-molecule imaging

PEGylated-biotin- and PEGylated-poly(T)-coated microscope slides were prepared based on the protocol described by Chandradoss et al ^49^. Ibidi glass coverslips (25 mm × 75 mm, IBIDI, IBD-10812) were cleaned with (1) MilliQ water (three washes, 5-min sonication, three washes), (2) 2% Alconox (Sigma, 242985; 20-min sonication followed by five washes with MilliQ water) and (3) 100% acetone (20-min sonication followed by three washes with MilliQ water). Slides were further cleaned and functionalized (hydroxylated) by incubation in 1 M KOH (Sigma, 484016) solution for 20 min while sonicating, followed by three washes with MilliQ water. Slides were sonicated twice for 10 min in 100% high-performance liquid chromatography (HPLC) ethanol (J.T. Baker, 8462-25) before applying amino-silanization chemistry. Next, slides were incubated for 24 min in a mixture of 3% 3-aminopropyltriethoxysilane (ACROS Organics, 430941000) and 5% acetic acid in HPLC ethanol, with a 1-min sonication in the middle. Slides were then washed with HPLC ethanol three times and MilliQ water three times and dried with nitrogen. Surface functionalization along with the first passivation step were performed by applying mPEG:

PEGylated-biotin/PEG-azide solution (20 mg PEGylated-biotin (Laysan, Biotin-PEG-SVA-5000) and 180 mg mPEG (Laysan, MPEG-SVA-5000) or 20 mg PEG-azide (JenKem, A5088) and 180 mg mPEG (Laysan, MPEG-SVA-5000)) dissolved in 1,560 µl of 0.1 M sodium bicarbonate (Sigma, S6297) and degassed (centrifugation for 1 min at 16,000g). Next, 140 µl of solution was applied on one surface, followed by immediate assembly of another surface on top. Each pair of assembled surfaces were incubated overnight in a dark, humid environment.

For PEGylated-biotin surfaces, the next day, surfaces were washed with MilliQ water and dried with nitrogen followed by a second passivation step. MS(PEG)4 (Thermo Fisher Scientific, TS-22341) was diluted in 0.1 M sodium bicarbonate to a final concentration of 11.7 mg ml–1 and applied on one surface, followed by the assembly of another surface on top. Each pair of assembled surfaces was incubated overnight in a dark, humid environment. The next day, surfaces were disassembled, washed with MilliQ water and dried with nitrogen. After nitrogen flush, surfaces were stored at –20 °C.

For PEGylated-poly(T) surfaces, following PEG-azide coating, surfaces were washed with MilliQ water and dried with nitrogen. To enable anchoring of dT50 to the surface via click chemistry, 10 µM 5′heyxynyl-dT50 (IDT) was mixed with 2 mM CuSO4 (Sigma, C1297) and double-distilled water. Next, 100 µl of the mixture was applied on one surface, followed by immediate assembly of another surface on top. Each pair of assembled surfaces was incubated overnight in a dark, humid environment. The next day, a second passivation step (MS(PEG)4) was performed similar to the PEGylated-biotin surface preparation. Finally, surfaces were stored at −20 °C after nitrogen flush.

### Antibody labeling

Capture and detection antibodies were labeled using a biotin conjugation kit (Abcam, ab201796) and Alexa Flour antibody-labeling kits (Thermo Fisher Scientific, A20181/A10237/A20186) according to the manufacturer’s protocol. Labeled antibodies were purified by size-exclusion chromatography using Bio-Spin 6 columns (Bio-Rad, 7326200) followed by measurement of protein concentration using a NanoDrop 2000 at 280 nm.

### Single-molecule imaging

PEGylated-biotin- and PEGylated-poly(T)-coated coverslips were assembled into an Ibidi flowcell (Sticky Slide VI hydrophobic, IBIDI, IBD-80608), generating a six-lane flowcell or ChipSop flowcell (chip 2 – 28 mini Luer platform Fl. 342, 30001180), generating a fourteen-lane flowcell, which enables imaging of six different samples or various combinations of antibodies on a single surface. For PEGylated-biotin flowcells, streptavidin (Sigma, S4762) was added to a final concentration of 0.2 mg/ml followed by a 10-min incubation and washing with imaging buffer (IMB; 12 mM HEPES pH 8 (Thermo Fisher Scientific, 15630056), 40 mM Tris pH 7.5 (Gibco, 115567-027), 60 mM KCl (Sigma, 60142), 0.32 mM EDTA (Invitrogen, 15575-038), 3 mM MgCl2 (Sigma, 63069), 10% glycerol (Bio-Lab, 56815), 0.1 mg/ml bovine serum albumin (BSA; Sigma, A7906) and 0.02% IGEPAL (Sigma, I8896)). For time-lapse imaging experiments (histone PTMs and p53), before sample application, TetraSpeck beads (Thermo Fisher Scientific, T7279) diluted in PBS were added and incubated on the surface for at least 10 min to allow correction for stage drift in image analysis. Imaging was performed on a TIRF microscope manufactured by Nikon (Eclipse Ti2-E LU-N4 TIRF) with a CFI apochromat TIRF ×60 objective lens and TRF49904, TRF49909 and TRF89902 filter cubes (Chroma) for the 488-, 561- and 647-nm lasers, respectively. Images were taken with the ×1.5 magnification setting, resulting in fields of view (FOVs) of 148 µm × 148 µm using an Andor Zyla 4.2 Plus camera. At least 50 FOVs were imaged per lane.

### Histone PTM analysis

PEGylated-poly(T)-coated coverslips were assembled as described and further passivated with 5% BSA (Merck, A7906) for 30 min, followed by washing with IMB. Next, plasma samples containing tailed and fluorescently labeled cfNuc were incubated with antibodies (diluted 1:60) for 30 min at room temperature to allow for the formation of antibody–cfNuc complexes. Next, samples were loaded on the surface and incubated for 15 min to allow hybridization. The flowcell was washed three times with IMB, followed by time-lapse imaging every 15 min with the three laser channels across all positions (50 FOVs, 148 µm2, per experiment).

### H3-K27M Enrichment analysis

PEGylated-biotin-coated coverslips were assembled and coated with streptavidin. Next, Biotinylated α-H3K27M antibody (Capture antibody) were incubated on the surface in IMB for 30 min, followed by 3 washes with IMB. Then, 45 ul of plasma sample was added to the flow cell and incubated on the surface for 30 min to allow binding of mutant histones, followed by three washes with IMB. Finally, fluorescently labeled α-H3K7Ac antibody (detection antibody) were introduced to the surface for 60 min, washed with IMB and imaged.

### Multiplexed p53 analysis

PEGylated-biotin-coated coverslips were assembled and coated with streptavidin. Biotinylated antibodies targeting all forms of p53 were incubated on the surface in IMB2 (10 mM MES pH 6.5 (Boston Bioproducts, NC9904354), 60 mM KCl, 0.32 mM EDTA, 3 mM MgCl2, 10% glycerol, 0.1 mg ml–1 BSA and 0.02% Igepal) for 30 min, followed by washing with IMB2. Next, the plasma sample was added to the flowcell and incubated on the surface for 30 min, followed by three washes with IMB2 to allow binding of target p53 proteins. Then, two distinct fluorescently labeled antibodies (detection antibodies); one antibody targeting both the wild-type and mutant conformation of p53 (total, red) and a second antibody which specifically recognize the mutant p53 conformation fluorescently were introduced to the surface for 10 min, followed by time-lapse imaging every 15 min with two laser channels across all positions (50 FOVs, 148 µm2, per experiment).

### Image analysis

Image analysis was performed with the open-source software CellProfiler v3.1.9 (http://www.cellprofiler.org/). Briefly, time-lapse images of antibody-binding events and TetraSpeck beads were aligned, stacked and summed to one image. Antibody spots can be differentiated from TetraSpeck bead spots based on spot size and intensity. Summed antibody images were aligned with cfNucleosome images to count colocalization events.

For oncohistones analysis, single time-point images of α-H3K27ac antibody binding events were quantified. To minimize surface batch effect, both H3-K27M-H3K27ac and p53 signals were normalized to the surface activity (average antibody signal across all surface.

### Machine learning model for sample classification

For sample binary classification, various machine learning algorithms were trained on the H3K27ac normalized score feature that showed significant differences between H3 WT and mutant DMG patients (figure 3C, D and Supplementary figure 3A) and evaluated for performance using a fourfold cross-validation across all samples. The best predictive performance was achieved by a Naïve Bayes classifier. To evaluate the classifier overall performance using the selected features, we performed repeated (10,000) fourfold cross-validation across all samples. For each iteration, the sensitivity, specificity, accuracy, precision, F1 score, and AUC value were calculated and averaged over all iterations. Scikit-learn and SciPy packages were used for machine learning modeling.

### Statistical analysis

All statistical analyses were conducted using the statistical programming language R v4.0.0. All tests of significance (Wilcoxon rank-sum exact test and Kruskal-Wallis H test) were conducted two sided. Multiple comparison corrections were calculated using Bonferroni Correction via the Rstatix package

## References

1. Sturm, D., Witt, H., Hovestadt, V., Khuong-Quang, D.A., Jones, D.T., Konermann, C., Pfaff, E., Tönjes, M., Sill, M., Bender, S., et al. (2012). Hotspot mutations in H3F3A and IDH1 define distinct epigenetic and biological subgroups of glioblastoma. Cancer Cell 22, 425–437. 10.1016/j.ccr.2012.08.024.

2. Karremann, M., Gielen, G.H., Hoffmann, M., Wiese, M., Colditz, N., Warmuth-Metz, M., Bison, B., Claviez, A., van Vuurden, D.G., von Bueren, A.O., et al. (2018). Diffuse high-grade gliomas with H3 K27M mutations carry a dismal prognosis independent of tumor location. Neuro Oncol 20, 123–131. 10.1093/neuonc/nox149.

3. Louis, D.N., Perry, A., Reifenberger, G., von Deimling, A., Figarella-Branger, D., Cavenee, W.K., Ohgaki, H., Wiestler, O.D., Kleihues, P., and Ellison, D.W. (2016). The 2016 World Health Organization Classification of Tumors of the Central Nervous System: a summary. Acta Neuropathol 131, 803–820. 10.1007/s00401-016-1545-1.

4. Louis, D.N., Perry, A., Wesseling, P., Brat, D.J., Cree, I.A., Figarella-Branger, D., Hawkins, C., Ng, H.K., Pfister, S.M., Reifenberger, G., et al. (2021). The 2021 WHO Classification of Tumors of the Central Nervous System: a summary. Neuro Oncol 23, 1231–1251. 10.1093/neuonc/noab106.

5. Hamisch, C., Kickingereder, P., Fischer, M., Simon, T., and Ruge, M.I. (2017). Update on the diagnostic value and safety of stereotactic biopsy for pediatric brainstem tumors: a systematic review and meta-analysis of 735 cases. J Neurosurg Pediatr 20, 261–268. 10.3171/2017.2.Peds1665.

6. Gupta, N., Goumnerova, L.C., Manley, P., Chi, S.N., Neuberg, D., Puligandla, M., Fangusaro, J., Goldman, S., Tomita, T., Alden, T., et al. (2018). Prospective feasibility and safety assessment of surgical biopsy for patients with newly diagnosed diffuse intrinsic pontine glioma. Neuro Oncol 20, 1547–1555. 10.1093/neuonc/noy070.

7. Van Mieghem, E., Wozniak, A., Geussens, Y., Menten, J., De Vleeschouwer, S., Van Calenbergh, F., Sciot, R., Van Gool, S., Bechter, O.E., Demaerel, P., et al. (2013). Defining pseudoprogression in glioblastoma multiforme. Eur J Neurol 20, 1335–1341. 10.1111/ene.12192.

8. Brandes, A.A., Tosoni, A., Spagnolli, F., Frezza, G., Leonardi, M., Calbucci, F., and Franceschi, E. (2008). Disease progression or pseudoprogression after concomitant radiochemotherapy treatment: pitfalls in neurooncology. Neuro Oncol 10, 361–367. 10.1215/15228517-2008-008.

9. Schwartzentruber, J., Korshunov, A., Liu, X.-Y., Jones, D.T.W., Pfaff, E., Jacob, K., Sturm, D., Fontebasso, A.M., Quang, D.-A.K., Tönjes, M., et al. (2012). Driver mutations in histone H3.3 and chromatin remodelling genes in paediatric glioblastoma. Nature 482, 226–231. 10.1038/nature10833.

10. Wu, G., Broniscer, A., McEachron, T.A., Lu, C., Paugh, B.S., Becksfort, J., Qu, C., Ding, L., Huether, R., Parker, M., et al. (2012). Somatic histone H3 alterations in pediatric diffuse intrinsic pontine gliomas and non-brainstem glioblastomas. Nat Genet 44, 251–253. 10.1038/ng.1102.

11. Bender, S., Tang, Y., Lindroth, Anders M., Hovestadt, V., Jones, David T.W., Kool, M., Zapatka, M., Northcott, Paul A., Sturm, D., Wang, W., et al. (2013). Reduced H3K27me3 and DNA Hypomethylation Are Major Drivers of Gene Expression in K27M Mutant Pediatric High-Grade Gliomas. Cancer Cell 24, 660–672. 10.1016/j.ccr.2013.10.006.

12. Mackay, A., Burford, A., Carvalho, D., Izquierdo, E., Fazal-Salom, J., Taylor, K.R., Bjerke, L., Clarke, M., Vinci, M., Nandhabalan, M., et al. (2017). Integrated Molecular Meta-Analysis of 1,000 Pediatric High-Grade and Diffuse Intrinsic Pontine Glioma. Cancer Cell 32, 520–537.e525. 10.1016/j.ccell.2017.08.017.

13. Harpaz, N., Mittelman, T., Beresh, O., Griess, O., Furth, N., Salame, T.-M., Oren, R., Fellus-Alyagor, L., Harmelin, A., Alexandrescu, S., et al. (2022). Single-cell epigenetic analysis reveals principles of chromatin states in H3.3-K27M gliomas. Molecular Cell 82, 2696–2713.e2699. 10.1016/j.molcel.2022.05.023.

14. Heitzer, E., Haque, I.S., Roberts, C.E.S., and Speicher, M.R. (2019). Current and future perspectives of liquid biopsies in genomics-driven oncology. Nat Rev Genet 20, 71–88. 10.1038/s41576-018-0071-5.

15. Bettegowda, C., Sausen, M., Leary, R.J., Kinde, I., Wang, Y., Agrawal, N., Bartlett, B.R., Wang, H., Luber, B., Alani, R.M., et al. (2014). Detection of circulating tumor DNA in early- and late-stage human malignancies. Sci Transl Med 6, 224ra224. 10.1126/scitranslmed.3007094.

16. Wadden, J., Ravi, K., John, V., Babila, C.M., and Koschmann, C. (2022). Cell-Free Tumor DNA (cf-tDNA) Liquid Biopsy: Current Methods and Use in Brain Tumor Immunotherapy. Front Immunol 13, 882452. 10.3389/fimmu.2022.882452.

17. Wang, Y., Springer, S., Zhang, M., McMahon, K.W., Kinde, I., Dobbyn, L., Ptak, J., Brem, H., Chaichana, K., Gallia, G.L., et al. (2015). Detection of tumor-derived DNA in cerebrospinal fluid of patients with primary tumors of the brain and spinal cord. Proc Natl Acad Sci U S A 112, 9704–9709. 10.1073/pnas.1511694112.

18. Miller, A.M., Shah, R.H., Pentsova, E.I., Pourmaleki, M., Briggs, S., Distefano, N., Zheng, Y., Skakodub, A., Mehta, S.A., Campos, C., et al. (2019). Tracking tumour evolution in glioma through liquid biopsies of cerebrospinal fluid. Nature 565, 654–658. 10.1038/s41586-019-0882-3.

19. Cantor, E., Wierzbicki, K., Tarapore, R.S., Ravi, K., Thomas, C., Cartaxo, R., Nand Yadav, V., Ravindran, R., Bruzek, A.K., Wadden, J., et al. (2022). Serial H3K27M cell-free tumor DNA (cf-tDNA) tracking predicts ONC201 treatment response and progression in diffuse midline glioma. Neuro Oncol 24, 1366–1374. 10.1093/neuonc/noac030.

20. Izquierdo, E., Proszek, P., Pericoli, G., Temelso, S., Clarke, M., Carvalho, D.M., Mackay, A., Marshall, L.V., Carceller, F., Hargrave, D., et al. (2021). Droplet digital PCR-based detection of circulating tumor DNA from pediatric high grade and diffuse midline glioma patients. Neurooncol Adv 3, vdab013. 10.1093/noajnl/vdab013.

21. Panditharatna, E., Kilburn, L.B., Aboian, M.S., Kambhampati, M., Gordish-Dressman, H., Magge, S.N., Gupta, N., Myseros, J.S., Hwang, E.I., Kline, C., et al. (2018). Clinically Relevant and Minimally Invasive Tumor Surveillance of Pediatric Diffuse Midline Gliomas Using Patient-Derived Liquid Biopsy. Clin Cancer Res 24, 5850–5859. 10.1158/1078-0432.Ccr-18-1345.

22. Bruzek, A.K., Ravi, K., Muruganand, A., Wadden, J., Babila, C.M., Cantor, E., Tunkle, L., Wierzbicki, K., Stallard, S., Dickson, R.P., et al. (2020). Electronic DNA Analysis of CSF Cell-free Tumor DNA to Quantify Multi-gene Molecular Response in Pediatric High-grade Glioma. Clin Cancer Res 26, 6266–6276. 10.1158/1078-0432.Ccr-20-2066.

23. Stallard, S., Savelieff, M.G., Wierzbicki, K., Mullan, B., Miklja, Z., Bruzek, A., Garcia, T., Siada, R., Anderson, B., Singer, B.H., et al. (2018). CSF H3F3A K27M circulating tumor DNA copy number quantifies tumor growth and in vitro treatment response. Acta Neuropathol Commun 6, 80. 10.1186/s40478-018-0580-7.

24. Sadeh, R., Sharkia, I., Fialkoff, G., Rahat, A., Gutin, J., Chappleboim, A., Nitzan, M., Fox-Fisher, I., Neiman, D., Meler, G., et al. (2021). ChIP-seq of plasma cell-free nucleosomes identifies gene expression programs of the cells of origin. Nat Biotechnol 39, 586–598. 10.1038/s41587-020-00775-6.

25. Gezer, U., Yörüker, E.E., Keskin, M., Kulle, C.B., Dharuman, Y., and Holdenrieder, S. (2015). Histone Methylation Marks on Circulating Nucleosomes as Novel Blood-Based Biomarker in Colorectal Cancer. Int J Mol Sci 16, 29654–29662. 10.3390/ijms161226180.

26. Snyder, M.W., Kircher, M., Hill, A.J., Daza, R.M., and Shendure, J. (2016). Cell-free DNA Comprises an In Vivo Nucleosome Footprint that Informs Its Tissues-Of-Origin. Cell 164, 57–68. 10.1016/j.cell.2015.11.050.

27. Sun, K., Jiang, P., Cheng, S.H., Cheng, T.H.T., Wong, J., Wong, V.W.S., Ng, S.S.M., Ma, B.B.Y., Leung, T.Y., Chan, S.L., et al. (2019). Orientation-aware plasma cell-free DNA fragmentation analysis in open chromatin regions informs tissue of origin. Genome Res 29, 418–427. 10.1101/gr.242719.118.

28. Van den Ackerveken, P., Lobbens, A., Turatsinze, J.-V., Solis-Mezarino, V., Völker-Albert, M., Imhof, A., and Herzog, M. (2021). A novel proteomics approach to epigenetic profiling of circulating nucleosomes. Scientific Reports 11, 7256. 10.1038/s41598-021-86630-3.

29. Fedyuk, V., Erez, N., Furth, N., Beresh, O., Andreishcheva, E., Shinde, A., Jones, D., Zakai, B.B., Mavor, Y., Peretz, T., et al. (2023). Multiplexed, single-molecule, epigenetic analysis of plasma-isolated nucleosomes for cancer diagnostics. Nature Biotechnology 41, 212–221. 10.1038/s41587-022-01447-3.

30. Furth, N., Algranati, D., Dassa, B., Beresh, O., Fedyuk, V., Morris, N., Kasper, L.H., Jones, D., Monje, M., Baker, S.J., and Shema, E. (2022). H3-K27M-mutant nucleosomes interact with MLL1 to shape the glioma epigenetic landscape. Cell Reports 39, 110836. 10.1016/j.celrep.2022.110836.

31. Harutyunyan, A.S., Chen, H., Lu, T., Horth, C., Nikbakht, H., Krug, B., Russo, C., Bareke, E., Marchione, D.M., Coradin, M., et al. (2020). H3K27M in Gliomas Causes a One-Step Decrease in H3K27 Methylation and Reduced Spreading within the Constraints of H3K36 Methylation. Cell Reports 33, 108390. 10.1016/j.celrep.2020.108390.

32. Lewis, P.W., Müller, M.M., Koletsky, M.S., Cordero, F., Lin, S., Banaszynski, L.A., Garcia, B.A., Muir, T.W., Becher, O.J., and Allis, C.D. (2013). Inhibition of PRC2 Activity by a Gain-of-Function H3 Mutation Found in Pediatric Glioblastoma. Science 340, 857–861. doi:10.1126/science.1232245.

33. Mohammad, F., Weissmann, S., Leblanc, B., Pandey, D.P., Højfeldt, J.W., Comet, I., Zheng, C., Johansen, J.V., Rapin, N., Porse, B.T., et al. (2017). EZH2 is a potential therapeutic target for H3K27M-mutant pediatric gliomas. Nature Medicine 23, 483–492. 10.1038/nm.4293.

34. Venneti, S., Kawakibi, A.R., Ji, S., Waszak, S.M., Sweha, S.R., Mota, M., Pun, M., Deogharkar, A., Chung, C., Tarapore, R.S., et al. (2023). Clinical Efficacy of ONC201 in H3K27M-Mutant Diffuse Midline Gliomas Is Driven by Disruption of Integrated Metabolic and Epigenetic Pathways. Cancer Discov 13, 2370–2393. 10.1158/2159-8290.Cd-23-0131.

35. Lo, Y.M.D., Han, D.S.C., Jiang, P., and Chiu, R.W.K. (2021). Epigenetics, fragmentomics, and topology of cell-free DNA in liquid biopsies. Science 372, eaaw3616. doi:10.1126/science.aaw3616.

36. Moss, J., Magenheim, J., Neiman, D., Zemmour, H., Loyfer, N., Korach, A., Samet, Y., Maoz, M., Druid, H., Arner, P., et al. (2018). Comprehensive human cell-type methylation atlas reveals origins of circulating cell-free DNA in health and disease. Nat Commun 9, 5068. 10.1038/s41467-018-07466-6.

37. Shen, S.Y., Singhania, R., Fehringer, G., Chakravarthy, A., Roehrl, M.H.A., Chadwick, D., Zuzarte, P.C., Borgida, A., Wang, T.T., Li, T., et al. (2018). Sensitive tumour detection and classification using plasma cell-free DNA methylomes. Nature 563, 579–583. 10.1038/s41586-018-0703-0.

38. Piunti, A., Hashizume, R., Morgan, M.A., Bartom, E.T., Horbinski, C.M., Marshall, S.A., Rendleman, E.J., Ma, Q., Takahashi, Y.-h., Woodfin, A.R., et al. (2017). Therapeutic targeting of polycomb and BET bromodomain proteins in diffuse intrinsic pontine gliomas. Nature Medicine 23, 493–500. 10.1038/nm.4296.

39. Stafford, J.M., Lee, C.H., Voigt, P., Descostes, N., Saldaña-Meyer, R., Yu, J.R., Leroy, G., Oksuz, O., Chapman, J.R., Suarez, F., et al. (2018). Multiple modes of PRC2 inhibition elicit global chromatin alterations in H3K27M pediatric glioma. Sci Adv 4, eaau5935. 10.1126/sciadv.aau5935.

40. Przystal, J.M., Cosentino, C.C., Yadavilli, S., Zhang, J., Laternser, S., Bonner, E.R., Biery, M., Vitanza, N.A., Koschmann, C., Cain, J., et al. (2021). HGG-32. ONC201 AND ONC206 TARGET TUMOR CELL METABOLISM IN PEDIATRIC DIFFUSE MIDLINE GLIOMA PRECLINICAL MODELS. Neuro-Oncology 23, i23–i24. 10.1093/neuonc/noab090.096.

41. Klein, E.A., Richards, D., Cohn, A., Tummala, M., Lapham, R., Cosgrove, D., Chung, G., Clement, J., Gao, J., Hunkapiller, N., et al. (2021). Clinical validation of a targeted methylation-based multi-cancer early detection test using an independent validation set. Ann Oncol 32, 1167–1177. 10.1016/j.annonc.2021.05.806.

42. Chen, X., Dong, Z., Hubbell, E., Kurtzman, K.N., Oxnard, G.R., Venn, O., Melton, C., Clarke, C.A., Shaknovich, R., Ma, T., et al. (2021). Prognostic Significance of Blood-Based Multi-cancer Detection in Plasma Cell-Free DNA. Clin Cancer Res 27, 4221–4229. 10.1158/1078-0432.Ccr-21-0417.

43. Mair, R., and Mouliere, F. (2022). Cell-free DNA technologies for the analysis of brain cancer. Br J Cancer 126, 371–378. 10.1038/s41416-021-01594-5.

44. Peneder, P., Stütz, A.M., Surdez, D., Krumbholz, M., Semper, S., Chicard, M., Sheffield, N.C., Pierron, G., Lapouble, E., Tötzl, M., et al. (2021). Multimodal analysis of cell-free DNA whole-genome sequencing for pediatric cancers with low mutational burden. Nat Commun 12, 3230. 10.1038/s41467-021-23445-w.

45. Mandal, S., Li, Z., Chatterjee, T., Khanna, K., Montoya, K., Dai, L., Petersen, C., Li, L., Tewari, M., Johnson-Buck, A., and Walter, N.G. (2021). Direct Kinetic Fingerprinting for High-Accuracy Single-Molecule Counting of Diverse Disease Biomarkers. Accounts of Chemical Research 54, 388–402. 10.1021/acs.accounts.0c00621.

46. Furth, N., Shilo, S., Cohen, N., Erez, N., Fedyuk, V., Schrager, A.M., Weinberger, A., Dror, A.A., Zigron, A., Shehadeh, M., et al. (2021). Unified platform for genetic and serological detection of COVID-19 with single-molecule technology. PLoS One 16, e0255096. 10.1371/journal.pone.0255096.

47. Ding, Z., Wang, N., Ji, N., and Chen, Z.S. (2022). Proteomics technologies for cancer liquid biopsies. Mol Cancer 21, 53. 10.1186/s12943-022-01526-8.

48. Müller Bark, J., Kulasinghe, A., Chua, B., Day, B.W., and Punyadeera, C. (2020). Circulating biomarkers in patients with glioblastoma. Br J Cancer 122, 295–305. 10.1038/s41416-019-0603-6.

49. Chandradoss, S.D., Haagsma, A.C., Lee, Y.K., Hwang, J.H., Nam, J.M., and Joo, C. (2014). Surface passivation for single-molecule protein studies. J Vis Exp. 10.3791/50549.

