## Supplemental figures and methods for "Single-molecule systems for detection and monitoring of plasma circulating nucleosomes and oncoproteins in Diffuse Midline Glioma"

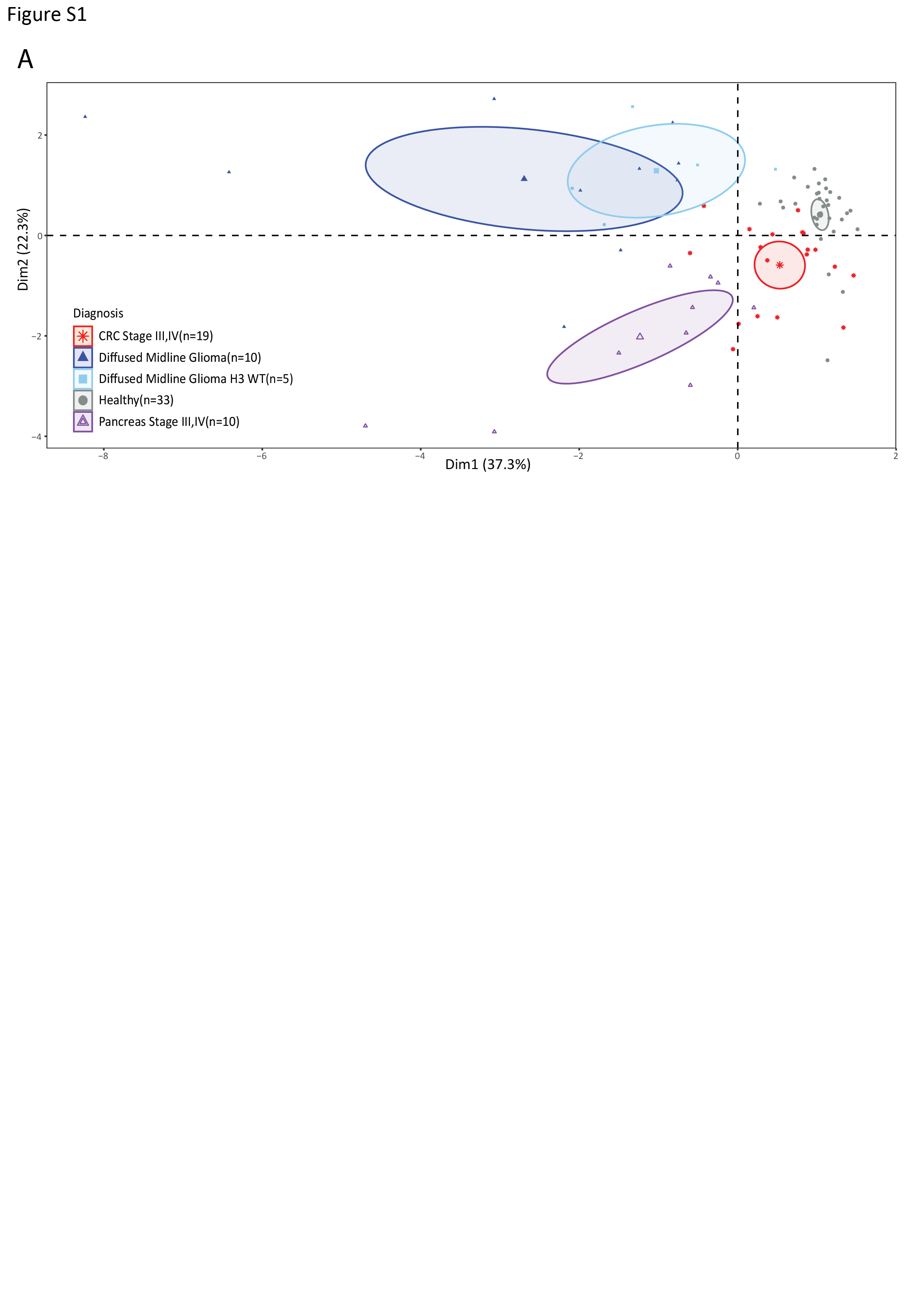

**Supplementary Figure 1**

**EPINUC analysis of plasma samples from DMG patients carrying WT H3 or H3-K27M-mutant, healthy individuals or patients diagnosed with CRC or PDAC.**

Principal Component Analysis (PCA), as presented in Fig.1C, noting the H3 status of the DMG samples. The following EPINUC parameters were used: H3K4me1/Nuc, H3K9me3/Nuc, H3K9ac/Nuc, H3K27me3/Nuc, H3K9ac/H3K4me1, H3K36me3/H3K9me3, H3K27me3 & H3K4me3, and Nucleosomes count. Sample groups are color-coded as indicated in the plot; each dot represents one plasma sample. Ellipse represents 95% confidence interval for the barycenter of each group (denoted with a larger symbol).

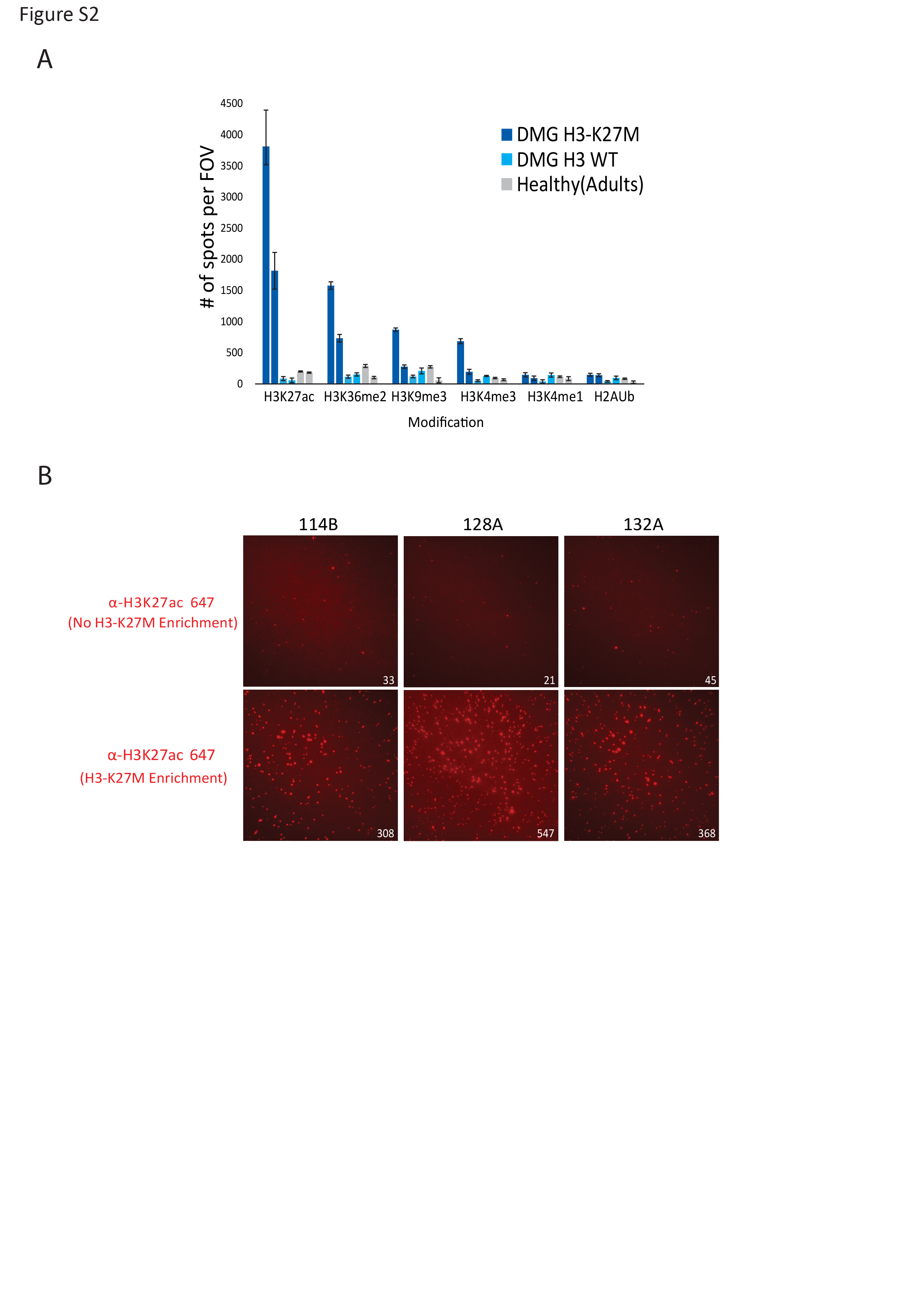

**Supplementary Figure 2**

**Single-molecule enrichment of plasma circulating H3-K27M Nucleosomes.**

**A.** Quantification of the single-molecule signal obtained for each tested antibody when incubated with H3-K27M enriched cfNuc from different plasma samples. Data is presented as the mean +/- s.d. of 50 FOVs per sample. **B.** Representative TIRF images of α-H3K27Ac signal of H3-K27M DMG plasma samples, originating from either all cfNuc (standard EPINUC analysis) or from on-surface enriched H3-K27M cfNuc. Identical samples were used for each comparison (sample ID is noted for each pair). Numbers within images represent counted spots; each spot correspond to a single surface bound nucleosome. Enriched H3-K27M cfNuc show higher H3K27ac signal

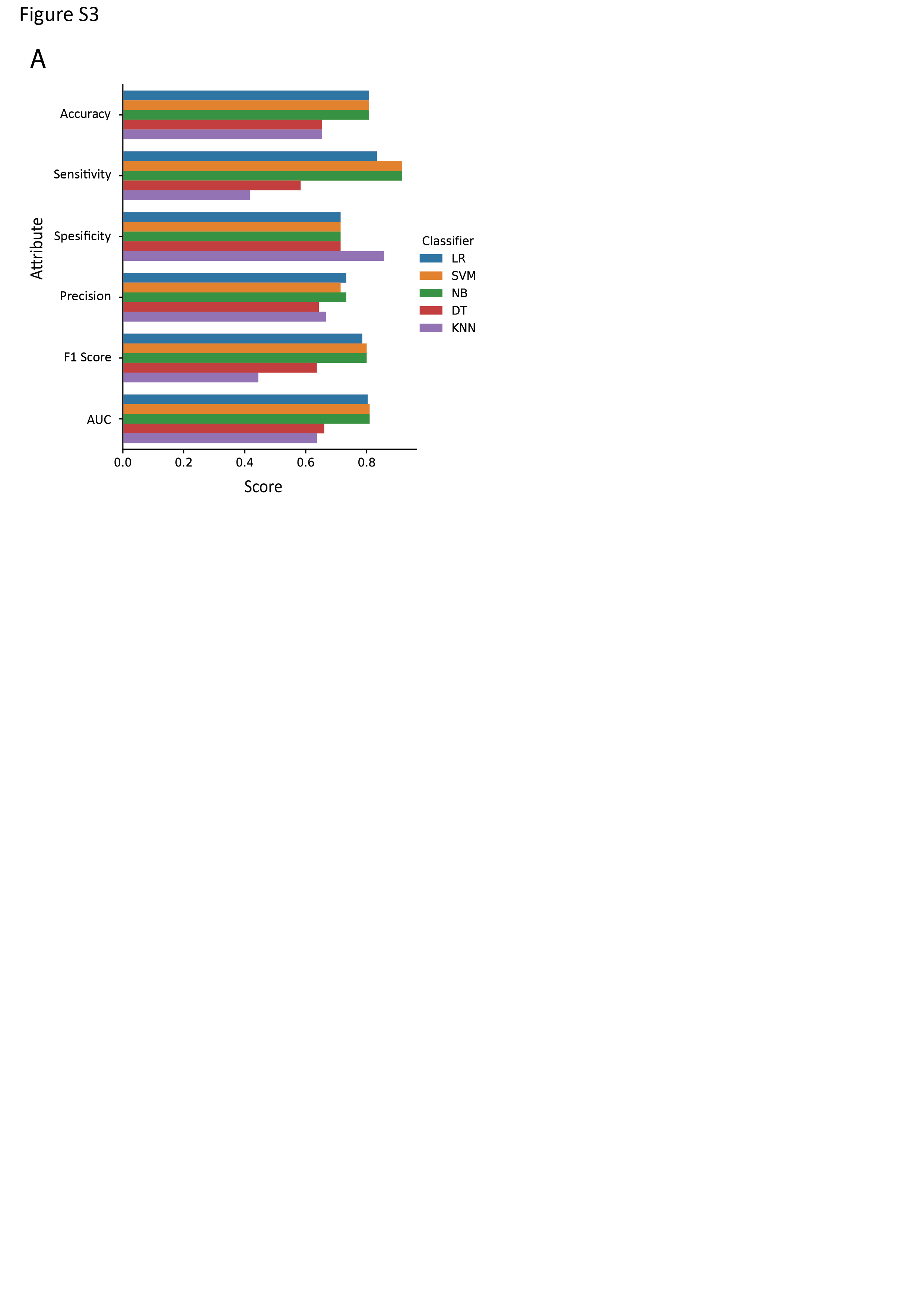

**Supplementary Figure 3**

**Prediction models performance comparison based on H3-K27M enrichment analysis.**

Machine learning algorithms performance comparison. Each bar presented as the mean value of 10,000 iterations and colored according to the corresponding model (Blue- Logistic Regression, Orange- Support Vector Machine, Green- Naïve Bayes, Red- Decision Tree, Purple- K Nearest Neighbors).

Detailed ddPCR primers, probes, and protocols

| Primer Name | Sequence |
| --- | --- |
| H3.3 K27M fwd | 5’ - CTCTGTACCATGGCTCGTA |
| H3.3 K27M rev | 5’ - CATACAAGAGAGACTTTGTCCC |
| H3.3 K27M Wt probe | /5HEX/TC+GC+A+A+GA+GT+GC /3IABkFQ/ |
| H3.3 K27M Mut probe | /5HEX/TC+G C+A+A +GA+G T+GC /3IABkFQ/ |
| TP53 R273C fwd | 5’ - GTAATCTACTGGGACGGAACAG |
| TP53 R273C rev | 5’ - CCCTTTCTTGCGGAGATTCT |
| TP53 R273C Wt probe | /5HEX/TT+GAGGT+G+C+GT+GTT /3IABkFQ/ |
| TP53 R273C Mut probe | /56-FAM/TG+AG+GT+G+T+GTGT/3IABkFQ/ |
| TP53 R175G fwd | 5’ - GCCATCTACAAGCAGTCACA |
| TP53 R175G rev | 5’ - CTGCTCACCATCGCTATCTG |
| TP53 R175G Wt probe | /5HEX/CAG+C+G+CCTC+ACA/3IABkFQ/ |
| TP53 R175G Mut probe | /56-FAM/CAG+C+C+CCTCAC/3IABkFQ/ |

‘+’ symbol indicates a locked nucleic acid and precedes the locked base. All primers and probes were sourced via Integrated DNA Technologies.

Pre-amplification protocol

After DNA extraction, elution volume was reduced to <20ul using a SpeedVac vacuum concentrator at 45ºC. Pre-amplification was performed in 50ul reactors using NEB Q5 Hot Start High-fidelity 2x Master Mix (NEB, M0494S) according to manufacturer’s instructions. Final primer concentrations for any singleplex or duplex amplifications were 0.5uM. Duplex pre-amplification protocol used for all targets is listed below:

| **Target** | **Initial Denature** | **Cycles** | **[Denature** | **Anneal** | **Extend]** | **Final Extension** |
| --- | --- | --- | --- | --- | --- | --- |
| H3.3 K27M + TP53 | 30s @ 98ºC | 35x | 10s @ 98ºC | 45s @ 58ºC | 10s @ 72C | 2min @72ºC |

After amplification, resulting duplex product was quantified, and diluted to ~100,000 copies/ul assuming 100% of resultant mass is made up of a 50/50 mix of H3.3 and TP53 amplicon product.

ddPCR protocol

Singleplex ddPCR reactions were prepared according to manufacturer’s instructions in 20ul reactors. Input template amount of 100,000 copies was used to calibrate the singleplex assays and adjusted if necessary depending on assay sensitivity requirements. Once calibrated, 3-6 technical replicates were performed for final analysis. Singleplex ddPCR amplification protocols used for all targets are listed below:

| **Target** | **Initial Denature** | **Cycles** | **[Denature** | **Probe Anneal** | **Extend]** | **Deactivation** |
| --- | --- | --- | --- | --- | --- | --- |
| H3.3 K27M | 10min @ 95ºC | 40x | 30s @ 94ºC | - | 1min @ 58C | 10min @98ºC |
| TP53 R273C | 10min @ 95ºC | 40x | 30s @ 94ºC | - | 1min @ 62C | 10min @98ºC |
| TP53 R175G | 10min @ 95ºC | 40x | 30s @ 94ºC | 30s @ 68C | 1min @ 60C | 10min @98ºC |
